# TurboID reveals the proxiomes of CGE1, VIPP1, and VIPP2 in *Chlamydomonas reinhardtii*

**DOI:** 10.1101/2022.12.01.518767

**Authors:** Elena Kreis, Katharina König, Frederik Sommer, Michael Schroda

## Abstract

In *Chlamydomonas reinhardtii*, VIPP1 and VIPP2 play a role in the sensing and coping with membrane stress and in thylakoid membrane biogenesis. To gain more insight into these processes, we aimed to identify proteins interacting with VIPP1/2 in the chloroplast and chose proximity labeling (PL) for this purpose. We used the transient interaction between the nucleotide exchange factor CGE1 and stromal HSP70B as test system. While PL with APEX2 and BioID proved to be inefficient, TurboID resulted in significant biotinylation *in vivo*. TurboID-mediated PL with VIPP1/2 as baits under ambient and H_2_O_2_ stress conditions confirmed known interactions of VIPP1 with VIPP2, HSP70B and CDJ2. Novel proteins in the VIPP1/2 interaction network can be grouped into proteins involved in the biogenesis of thylakoid membrane complexes and the regulation of photosynthetic electron transport. A third group comprises 11 proteins of unknown function whose genes are upregulated under chloroplast stress conditions. We named them VIPP PROXIMITY LABELING (VPL1-11). and confirmed the proximity of VIPP1 and VPL2 in a reciprocal experiment. Our results demonstrate the robustness of TurboID-mediated PL for studying protein interaction networks in the chloroplast of *Chlamydomonas* and pave the way for analyzing functions of VIPPs in thylakoid biogenesis and stress responses.

## Introduction

The identification of protein-protein interactions is key for the understanding of protein function. Traditional methods include affinity-purification combined with mass spectrometry (AP-MS) and derivatives like single- or tandem-affinity purification, pull-downs using immobilized bait proteins, co-migration on native gels (complexome profiling), yeast-two-hybrid and its derivatives. The disadvantage of all these methods is that protein interactions must persist during cell lysis and unspecific interactions may form after the mixing of compartment contents. Moreover, interactions are studied outside of the native context where important co-factors or scaffolds may be missing. Many of these limitations can be overcome by proximity labeling (PL) (reviewed in (Kim and Roux, 2016; Mair and Bergmann, 2021; Qin et al., 2021; Zhang et al., 2022)).

PL is based on the fusion of a bait to an enzyme that produces activated biotin (Roux et al., 2012). Activated biotin binds to amino acid side chains of proteins in the vicinity of the bait, which can be enriched with streptavidin beads and identified by mass spectrometry, thereby detecting stable as well as transient interactors and proteins in close proximity. PL is most commonly realized with two enzymes, BirA and APEX. The *E. coli* bifunctional ligase/repressor BirA activates biotin by adenylation, consuming ATP. Reactive biotinoyl-5’-AMP is normally transferred to a specific lysine residue of a subunit of the acetyl-CoA carboxylase requiring biotin as cofactor. This specificity is lost in BirA* harboring the R118G mutation such that reactive biotinyl-AMP is released and proteins in the vicinity of BirA* get biotinylated at exposed primary amines within an estimated range of approximately 10 nm (Choi-Rhee et al., 2004; Kim et al., 2014). Adding biotin to human cells expressing a fusion of BirA* with lamin-A in the nuclear lamina allowed the identification of novel interaction partners *in vivo* and the approach was coined BioID (proximity-dependent <u>bio</u>tin <u>id</u>entification) (Roux et al., 2012). The disadvantage of BirA* is its slow labeling kinetics, requiring labeling times of 15-18 h (Choi-Rhee et al., 2004; Roux et al., 2012). Moreover, BirA* is not very active at temperatures below 37°C (Kim et al., 2016; Zhang et al., 2019; Arora et al., 2020).

APEX (enhanced ascorbate peroxidase) is an engineered cytosolic peroxidase from pea or soybean originally destined for electron microscopy studies (Martell et al., 2012). If biotin-phenol and H_2_O_2_ are supplied to cells expressing APEX, biotin-phenol is converted into a biotin-phenoxyl radical that attacks electron-rich amino acid side chains of proximal proteins with a labeling radius of <20 nm. This approach was first used to map the mitochondrial matrix proteome in human cells (Rhee et al., 2013). APEX was then subjected to directed evolution yielding the much more active APEX2 (Lam et al., 2015). The advantage of APEX2 is the much faster labeling kinetics, requiring labeling times of less than a minute. The disadvantage is that biotin-phenol is less membrane permeable than biotin and that toxic H_2_O_2_ must be applied (Hwang and Espenshade, 2016; Tan et al., 2020).

The slow labeling kinetics of BirA* has been overcome recently by subjecting BirA* to directed evolution, resulting in TurboID and miniTurboID (Branon et al., 2018). TurboID achieves the same biotin labeling efficiency in 10 min as BirA* in 18 h, has greater activity at ambient temperatures and a slightly bigger labeling radius of ∼35 nm (May et al., 2020). Disadvantages of TurboID are baseline activity in the presence of endogenous biotin and inability to control enzyme activity by activation as it is possible in the APEX system. Nevertheless, since its publication in 2018, TurboID has been used extensively to map proteomes and to identify protein-interaction networks in a variety of model organisms including mammalian cells (Cho et al., 2020), zebrafish (Xiong et al., 2021), or yeast (Larochelle et al., 2019). There are also first reports on the successful application of TurboID to land plants for the identification of interactors of plant immune receptor N (Zhang et al., 2019), stomatal transcription factor FAMA (Mair et al., 2019), nuclear transport receptor exportin 4 (Xu et al., 2021), nuclear pore complex protein GBPL3 (Tang et al., 2022), and the TPLATE complex (Arora et al., 2020) as well as for the mapping of the nuclear proteome (Mair et al., 2019).

VESICLE-INDUCING PROTEIN IN PLASTIDS 1 (VIPP1) is a member of the ancient ESCRT-III membrane-remodelling superfamily. Cyanobacterial VIPP1 forms large basket-like assemblies (Gupta et al., 2021; Liu et al., 2021). Inside of the basket, N-terminal amphipathic α-helices (AH) of 24 amino acids length from each monomer align to form large hydrophobic columns, enabling the basket to bind to membranes and to encapsulate a vesicle-like bud. In the chloroplast of *Chlamydomonas reinhardtii* (*Chlamydomonas* hereafter), VIPP1 was found in long rods that tubulate membranes and in short rods that form connections between the inner envelope and thylakoids (Gupta et al., 2021). These connections could mediate lipid transfer between the inner envelope and thylakoids, explaining why *vipp1* knockout mutants of Arabidopsis have a severely reduced thylakoid membrane system (Zhang et al., 2012).

The chloroplast HSP70 chaperone system in *Chlamydomonas* consists of HEAT SHOCK PROTEIN 70B (HSP70B), nucleotide exchange factor CHLOROPLAST GRPE 1 (CGE1), and at least six J-domain proteins CHLOROPLAST DNAJ (CDJ1-6) (reviewed in Trösch et al. (2015)). The latter supply HSP70B with specific substrates, which are misfolded proteins in the case of CDJ1 (Willmund et al., 2008) and VIPP1 in the case of CDJ2 (Liu et al., 2005). HSP70B/CDJ2/CGE1 catalyze the ATP-dependent assembly of VIPP1 monomers/dimers into large assemblies and their disassembly to monomers/dimers *in vitro*. Large rods were completely disassembled by HSP70B/CDJ2/CGE1 *in vitro* (Liu et al., 2007).

Several lines of evidence suggest a role of VIPP1 in the biogenesis of thylakoid membrane protein complexes. First, VIPP1 improved protein export via the bacterial and thylakoidal twin-arginine transport pathways (DeLisa et al., 2004; Lo and Theg, 2012). Second, in an *in vitro* reconstituted system to study the co-translational insertion of the D1 protein into thylakoid membranes, VIPP1 stimulated the formation of a D1 insertion intermediate and was found complexed with cpSecY, Alb3 and cpFtsY (Walter et al., 2015). Third, Arabidopsis, *Chlamydomonas* and cyanobacterial *vipp1* knockdown mutants displayed reduced levels of at least one of the major thylakoid membrane protein complexes (Kroll et al., 2001; Fuhrmann et al., 2009; Nordhues et al., 2012; Zhang et al., 2014; Zhang et al., 2016a).

Another role of VIPP1 is related to the protection of chloroplast membranes from various stresses. *Chlamydomonas vipp1* knockdown mutants or cyanobacterial mutants with mutations in the AH showed swollen thylakoids after exposure to high light (Nordhues et al., 2012; Gupta et al., 2021), and mesophyll cells in Arabidopsis *vipp1* knockdown mutants exhibited swollen chloroplasts due to impaired envelope response to hypotonic membrane stress (Zhang et al., 2012). Moreover, VIPP1 overexpression enhanced the recovery of photosynthetic capacity after heat stress, which was proposed to be due to a membrane protective role of VIPP1 (Zhang et al., 2016b). And VIPP1 overexpression trans-complemented a chloroplast swelling phenotype in the Arabidopsis *ncy1* stay-green mutant proposed to result from oxidative membrane damage (Zhang et al., 2016a).

The mechanism by which VIPP1 protects damaged membranes was proposed to be related to its ability to insert the AH into membranes exhibiting stored curvature elastic stress (SCE). VIPP1 stabilizes stressed membranes by multiple AH insertions, alleviating SCE stress and imparting a scaffold effect that prevents the membrane phase transition into a porous state (McDonald et al., 2015; McDonald et al., 2017). An important determinant here is the hydrophobicity of the hydrophobic face of the AH. *Chlamydomonas* has a paralog of VIPP1 named VIPP2 (Nordhues et al., 2012). VIPP2 has a more hydrophobic AH than VIPP1 and binds more strongly to chloroplast membranes than VIPP1 in cells subjected to H_2_O_2_ stress (Theis et al., 2020). VIPP2 is barely expressed under ambient conditions but accumulates strongly under various stress conditions, including high light intensities or elevated cellular H_2_O_2_ concentrations (Nordhues et al., 2012; Blaby et al., 2015; Perlaza et al., 2019; Theis et al., 2020), the depletion of the ClpP protease or of thylakoid membrane transporters/integrases (Ramundo et al., 2014; Theis et al., 2020), the addition of nickel ions (Blaby-Haas et al., 2016), the addition of alkylating agents (Fauser et al., 2022), or the inhibition of chloroplast fatty acid synthesis (Heredia-Martínez et al., 2018). In addition to VIPP2, these stresses also result in the accumulation of chloroplast chaperones and proteases including CLPB3, HSP70B, HSP22E/F, and DEG1C, linking chloroplast membrane stress with protein homeostasis. Apparently, misfolded, misassembled, and aggregated proteins in chloroplast membranes cause SCE stress that must be coped with by coordinated action of VIPPs, chaperones, and proteases as part of a chloroplast-specific unfolded protein response (cpUPR) (McDonald et al., 2015; Theis et al., 2020). Accordingly, VIPP1, VIPP2, HSP70B, and HSP22E/F have been found to interact at stressed chloroplast membranes (Theis et al., 2020). The ability of the VIPPs to sense SCE stress could also be the source of a retrograde signal for the cpUPR, which is supported by the finding that the induction of *HSP22E/F* gene expression upon high light exposure was strongly impaired in the *vipp2* mutant (Theis et al., 2020).

The aim of this work was to establish PL in the *Chlamydomonas* chloroplast based on the transient interaction between CGE1 and HSP70B as test system with the eventual goal to identify the proxiomes of VIPP1 and VIPP2. We show that TurboID fused to CGE1, VIPP1, and VIPP2 resulted in efficient *in vivo* PL under ambient conditions, heat stress, and oxidative stress. Many known interactions were confirmed, underpinning the strength of PL for revealing protein-interaction networks. We identified CGE2 as a novel co-chaperone of CGE1/HSP70B. In the VIPP1/2 proxiomes generated in three experimental setups, we identified 11 proteins of unknown function whose encoding genes were upregulated under chloroplast stresses; one of which was validated via reciprocal PL. Other VIPP1/2 proxiome proteins play roles in the biogenesis of thylakoid membrane protein complexes and the regulation of photosynthetic electron transport.

## Results

### Proximity labeling (PL) with APEX2 in *Chlamydomonas* only works *in vitro*

To establish *in vivo* PL in the chloroplast of *Chlamydomonas*, we synthesized the sequence encoding engineered soybean ascorbate peroxidase APEX2 (Lam et al., 2015) as a level 0 standard gene part for the *Chlamydomonas* Modular Cloning (MoClo) kit (Crozet et al., 2018). To enhance gene expression, the sequence was codon-optimized and interrupted by the first *Chlamydomonas RBCS2* intron (Baier et al., 2018; Schroda, 2019). Moreover, the coding sequence of the chloroplast transit peptide of *Chlamydomonas* HSP70B (cp70B), including the first *HSP70B* intron, was added (Drzymalla et al., 1996) (Supplemental Figure S1). We chose the CGE1 nucleotide exchange factor of chloroplast HSP70B as bait, since it interacts only transiently with HSP70B in the ADP-bound state (Schroda et al., 2001; Schmollinger et al., 2012) and therefore can be regarded as a suitable test system for a transient *in vivo* interaction. Here we went for an N-terminal fusion of APEX2 to CGE1, as the N-termini of GrpE-type nucleotide exchange factors represent unstructured regions located proximal to the substrate-binding domain of their Hsp70 partners and could potentially bring APEX2 close to HSP70B substrates (Rosenzweig et al., 2019). We amplified the *CGE1* gene without sequences encoding the chloroplast transit peptide from genomic DNA and cloned it into a level 0 MoClo vector. Parts coding for cp70B-APEX2, CGE1, mCherry (as control), and a 3xHA tag were then assembled with the *HSP70A-RBCS2* fusion promoter and the *RPL23* terminator into transcription units (level 1) (Crozet et al., 2018). After adding the *aadA* spectinomycin resistance cassette, resulting level 2 constructs (Figure 1A; Supplemental Figure S2) were transformed into the UVM4 expression strain (Neupert et al., 2009). The APEX2-CGE1 fusion protein was expressed at high frequency and accumulated at higher levels than native CGE1 (Supplemental Figure S2A). The APEX2-mCherry fusion was also well expressed (Supplemental Figure S2B).

**Figure 1.**
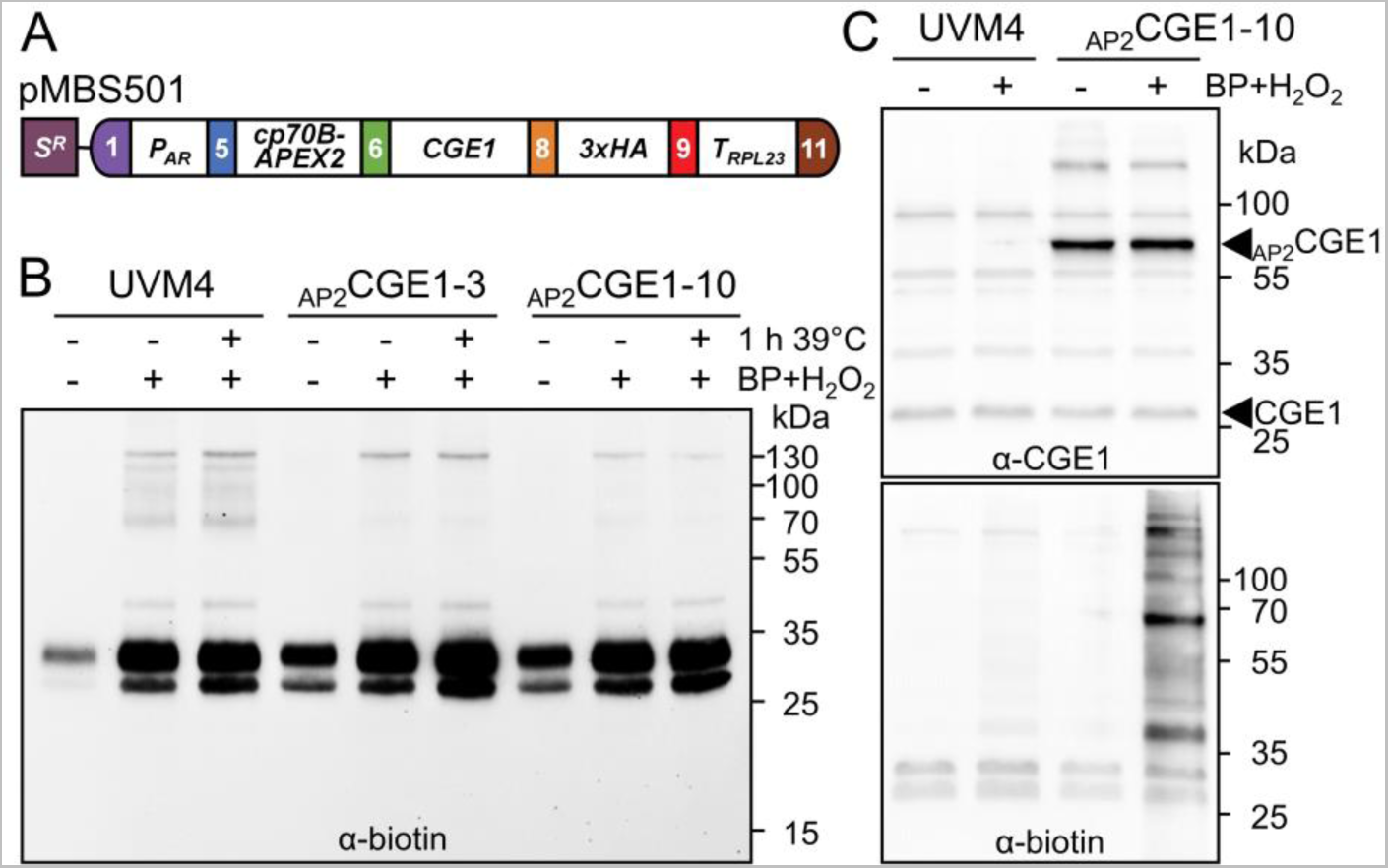
Biotin labeling with APEX2. **(A)** Level 2 construct pMBS501 conferring resistance to spectinomycin (S^R^) and driving the production of CGE1 fused N-terminally to APEX2 (_AP2_CGE1) and C-terminally to a 3xHA tag. Gene expression is controlled by the *HSP70A-RBCS2* promoter (*P_AR_*) and the *RPL23* terminator (*T_RPL23_*). The fusion protein is targeted to the chloroplast via the HSP70B chloroplast transit peptide (cp70B). (**B**) *In-vivo* labeling with biotin-phenol. *Chlamydomonas* cultures of the recipient strain UVM4 and two _AP2_CGE1 transformants (Supplemental Figure S2) were grown at 22°C to mid-log phase. One third of each culture was exposed to 39°C for 1 h. 500 µM biotin-phenol (BP) was added to the stressed and to half of the non-stressed culture for 30 min followed by the addition of 1 mM H_2_O_2_ for 1 min. Protein biotinylation in whole-cell proteins was analyzed by SDS-PAGE and immunoblotting using an antibody against biotin. **(C)** *In-vitro* labeling with biotin-phenol. *Chlamydomonas* cultures of the recipient strain UVM4 and an _AP2_CGE1 transformant were grown at 22°C to mid-log phase, harvested, and cells lysed by repeated freeze-thaw cycles. Soluble proteins were left untreated (-) or incubated with 500 µM BP for 30 min followed by the addition of 1 mM H_2_O_2_ for 1 min. Proteins were analyzed by SDS-PAGE and immunoblotting using antibodies against CGE1 (top) or biotin (bottom).

We then supplemented cultures of two lines expressing APEX2-CGE1 and the UVM4 control strain with 500 µM biotin-phenol and 1 mM H_2_O_2_ for 1 min to start APEX2-mediated biotinylation (Lam et al., 2015). However, an antibody against biotin did not detect any protein biotinylation in total protein extracts of APEX2-CGE1 producing strains in addition to endogenously biotinylated proteins also detected in the UVM4 control (Figure 1B). This was also observed when cells were exposed to 1 h of heat shock at 39°C prior to biotin-phenol and H_2_O_2_ addition. Specific biotinylation was not observed in APEX2-CGE1 and APEX2-mCherry production lines even when the preincubation with biotin-phenol was extended up to 24 hours or the reaction time with H_2_O_2_ was increased up to 1 hour (Supplemental Figure S3). In contrast, when biotin-phenol and H_2_O_2_ were added to soluble protein extracts, we detected several biotinylated proteins in an APEX2-CGE1 producing strain that were absent in the UVM4 control, with the most prominent biotinylated protein band migrating at the molecular mass of the APEX2-CGE1 fusion protein (Figure 1C). We conclude that APEX2-mediated PL only works *in vitro*, presumably because biotin-phenol is not taken up by intact *Chlamydomonas* cells.

### PL with TurboID in *Chlamydomonas* works *in vivo* and can be boosted by biotin addition

Since APEX2 did not support *in vivo* PL, we turned to the BioID (Roux et al., 2012) and TurboID systems (Branon et al., 2018). Codon-optimized DNA sequences coding for both proteins, interrupted by the 5^th^ *HSP70B* intron, were synthesized and sequences coding for the HSP70B chloroplast transit peptide, including the first *HSP70B* intron, were added to generate MoClo level 0 parts for N-terminal fusions. For TurboID, we also generated a level 0 part for C-terminal fusions (Supplemental Figure S1). Following the design used for APEX2 constructs, we assembled level 2 constructs for the production of BioID-CGE1 and TurboID-CGE1 (N-terminal fusions). Moreover, we assembled constructs for the production of VIPP1, VIPP2, and mCherry with C-terminal TurboID fusions (Figure 2A). VIPP1 fused C-terminally to GFP fully rescued the severe albino phenotype of *vipp1* knock-out mutants, suggesting that VIPP function is not impaired by such fusions (Zhang et al., 2012). Chloroplast-targeting of mCherry-TurboID was realized by the CDJ1 chloroplast transit peptide, while for VIPP1 and VIPP2 their native targeting peptides were employed.

**Figure 2.**
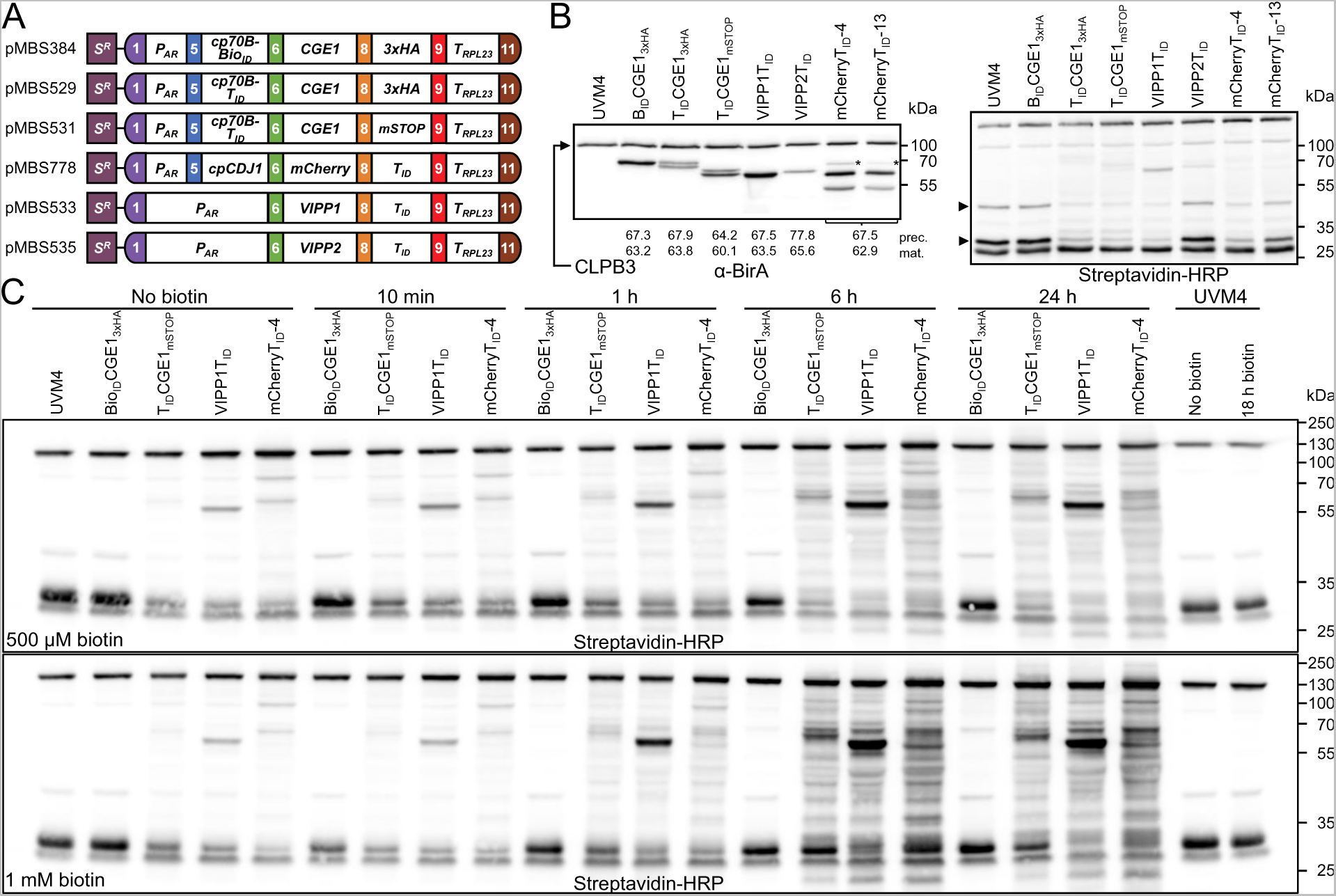
Construction of strains producing bait proteins fused with BioID and TurboID and analysis of their biotinylation patterns. **(A)** Level 2 constructs conferring resistance to spectinomycin (*S^R^*) and driving the production of bait proteins fused to BioID and TurboID as described in Figure 1A. CGE1 is produced with an N-terminal fusion to BioID (B_ID_) and TurboID (T_ID_) and targeted to the chloroplast via an HSP70B chloroplast transit peptide (cp70B). All other baits contain a C-terminal TurboID fusion. mCherry is targeted to the chloroplast via the chloroplast transit peptide of CDJ1 (cpCDJ1), VIPP1 and VIPP2 via their native transit peptides. (B) Analysis of fusion protein accumulation and self-biotinylation. Total cell protein extracts corresponding to 2 μg chlorophyll of the UVM4 recipient strain and transformants producing the four baits fused to BioID or TurboID were separated by SDS-PAGE and analyzed by immunoblotting using antibodies against BirA (left panel) or streptavidin coupled with horseradish peroxidase (HRP) (right panel) to capture biotinylated proteins. The home-made BirA antibody also detects plastidic CLPB3 (Supplemental Figure S6). Asterisks indicate putative chloroplast import precursors. Arrowheads point to proteins whose native biotinylation level decreases upon expression of TurboID fusions. Expected masses for precursor (prec) and mature (mat) proteins are given (Supplemental Table S3). Notice that GrpE proteins generally migrate with larger apparent than calculated masses (Schroda et al., 2001). (**C**) Enhancement of BioID and TurboID *in-vivo* biotinylation activity by externally added biotin. Mid-log phase cultures of the UVM4 recipient strains and transformants producing BioID/TurboID fusions with CGE1, VIPP1, and mCherry were supplied with 500 µM or 1 mM biotin and grown for the indicated times at 22°C. Protein biotinylation was analyzed by immunoblotting of total protein extracts corresponding to 2 µg chlorophyll using streptavidin-HRP.

BioID-CGE1 and TurboID-CGE1 accumulated to roughly the same levels as native CGE1 (Supplemental Figures S4 and S5A). BioID-CGE1 gave rise to a single protein band in SDS-PAGE, while TurboID-CGE1 gave rise to a double band (Figure 2B; Supplemental Figures S4 and S5A). Here it is not clear, whether the double band originates from alternative splicing of the *CGE1* transcript (Schroda et al., 2001) or from inefficient targeting to the chloroplast by the HSP70B transit peptide. Alternative splicing results in an additional Val-Gln dipeptide in the CGE1b isoform that can readily be resolved in SDS-PAGE and might well account for the small difference in apparent molecular mass observed (Willmund et al., 2007). In contrast, the larger size difference between the two bands observed for mCherry-TurboID (Figure 2B; Supplemental Figure S5B) suggests inefficient chloroplast targeting via the CDJ1 transit peptide, albeit this sequence did allow for efficient chloroplast-targeting of a heterologous cargo (Niemeyer et al., 2021). This might be due to the requirement of specific sequence stretches past the cleavage site for most *Chlamydomonas* chloroplast transit peptides, thereby rendering efficient chloroplast-targeting cargo-dependent (Caspari, 2022). The single bands detected for VIPP1- and VIPP2-TurboID imply efficient chloroplast targeting with the native transit peptides (Figure 2B). VIPP1-TurboID accumulated to much higher levels than VIPP2-TurboID, which was the most weakly expressed of all fusion proteins (Supplemental Figure S5C). Notice that VIPP2 is barely expressed under non-stress conditions (Nordhues et al., 2012; Theis et al., 2020).

We could detect constitutive *in-vivo* self-biotinylation (or cis-biotinylation, Arora et al. (2020)) without biotin addition for all TurboID fusion proteins but not for BioID-CGE1 despite its strong expression levels (Figure 2B). Interestingly, biotinylation levels of two naturally biotinylated proteins declined in cells expressing TurboID fusion proteins but not in cells expressing BioID-CGE1 (arrowheads in Figure 2B). The extent of this decline correlated with expression levels of TurboID fusions. Apparently, TurboID competes with the natural biotinylation machinery in the chloroplast for endogenous biotin. We wondered, whether a reduced abundance of naturally biotinylated proteins and the constitutive biotinylation activity of TurboID might affect chloroplast protein homeostasis and cellular fitness. To test this, we exposed lines producing TurboID-CGE1 to 40°C for 24 h and lines producing VIPP1/2-TurboID to high light of 1000 µE m^-2^ s^-1^ for 10 h, allowed them to recover for 16-24 h at 22°C, and analyzed levels of cpUPR markers VIPP1, CLPB3, HSP22E/F, HSP70B, and DEG1C (Ramundo et al., 2014; Perlaza et al., 2019). We found no differences in growth behavior or in the abundances of the cpUPR markers between the TurboID lines and the UVM4 control (Supplemental Figure S7). This suggests that the constitutive biotinylation activity of TurboID has no adverse effects, at least not under the conditions tested here.

Next we tested, whether *in-vivo* biotinylation via BioID and TurboID can be boosted by exogenously added biotin. To this end, we added 500 µM and 1 mM biotin to cultures of the UVM4 control and strains producing BioID- and TurboID fusions with CGE1, VIPP1, and mCherry. As shown in Figure 2C, with both concentrations of biotin we observed slightly increased cis- and trans-biotinylation already 10 min after biotin addition, which grew dramatically stronger 6 h after biotin addition, while no further biotinylation was observed between 6 h and 24 h after biotin addition. Protein biotinylation 1 h, 6 h and 24 h after biotin addition was higher when 1 mM biotin was added than when 500 µM biotin was added. Hence, addition of 1 mM biotin for 1 to 6 h appears optimal for boosting biotinylation of chloroplast proteins. Notice that weak cis-biotinylation of BioID-CGE1 became detectable only when biotin was added for at least 6 h.

We conclude that TurboID fused N- or C-terminally to different bait proteins allowed for efficient *in-vivo* protein biotinylation which can be boosted by the addition of biotin to the cultures. BioID does not promote efficient biotinylation. Naturally occurring biotinylation suffers proportional to TurboID expression levels. Chloroplast transit peptides cannot efficiently target any heterologous cargo to the organelle.

### Proof of concept: TurboID-CGE1 allows capturing the interaction with HSP70B without biotin boost and reveals CGE2 as novel co-chaperone

The efficient *in vivo*-biotinylation observed for TurboID-fusion proteins encouraged us to test, whether TurboID-mediated PL would allow capturing the transient interaction between CGE1 and HSP70B. To this end, we used a strain producing TurboID-CGE1 and, as controls, two strains producing mCherry-TurboID at different expression levels as well as the UVM4 recipient strain. Cells grown at ambient temperatures and heat-stressed at 40°C for 1 h were lysed and lysates incubated with streptavidin beads. We first analyzed proteins in the input and the streptavidin eluate by immunoblotting using streptavidin-HRP and antibodies against CGE1 and HSP70B, and known HSP70B (co-)chaperone partners HSP90C (Willmund and Schroda, 2005), CDJ1 (Willmund et al., 2008), and HSP22E/F (Rütgers et al., 2017) as well as the known HSP70B substrate VIPP1 (Liu et al., 2005). As shown in Figure 3A, naturally biotinylated proteins and proteins biotinylated via TurboID were clearly enriched with the streptavidin beads. TurboID-CGE1 was also enriched, as was native CGE1, pointing to cis-biotinylation of the fusion protein and its ability to interact with (and trans-biotinylate) native CGE1. HSP70B and its main J-domain co-chaperone CDJ1 were clearly enriched in the TurboID-CGE1 producing line versus the mCherry-TurboID and UVM4 controls. This enrichment was independent of whether cells were exposed to heat shock prior to lysis. No specific enrichment was found for HSP90C, HSP22E/F, and VIPP1. Rather, HSP22E/F after induction by heat shock appeared to interact unspecifically with the beads.

**Figure 3.**
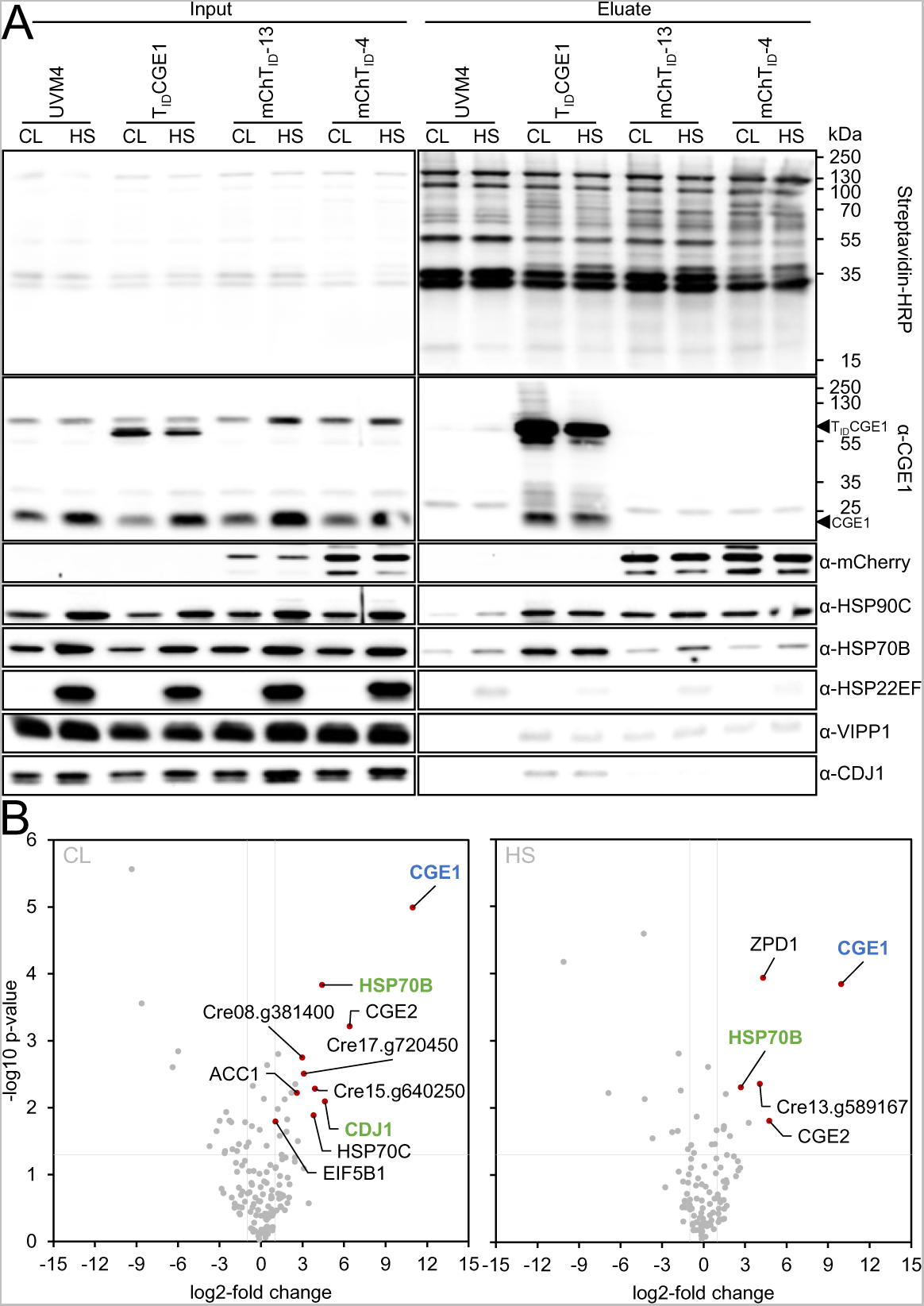
TurboID-based *in-vivo* proximity labeling without biotin boost using CGE1 as a bait. (**A**) Immunoblot analysis. Cultures of the UVM4 recipient strain, a transformant producing T_ID_CGE1, and two transformants accumulating different levels of mCherryT_ID_ were grown to mid-log phase at 22°C (CL). Half of the cultures was exposed to 40°C for 60 min (HS). Cells were harvested, lysed and biotinylated proteins were affinity-purified with streptavidin beads. Protein extracts before incubation (0,03% of the input) and after elution from the beads (5% of the eluate) were separated by SDS-PAGE and analyzed by immunoblotting with streptavidin-HRP or antibodies against the proteins indicated. The positions of the TurboID-CGE1 fusion protein (T_ID_CGE1) and of native CGE1 are indicated. One of three biological replicates is shown. **(B)** Volcano plots of the CGE1 proxiomes under ambient (CL) and heat stress (HS) conditions. Streptavidin bead eluates of the samples shown in (A) were analyzed by mass spectrometry. Shown is a comparison between protein abundances of the T_ID_CGE1 samples and two T_ID_mCherry lines after subtraction of contaminants and endogenous biotinylated proteins from the UVM4 control strain. Proteins significantly enriched in the T_ID_CGE1 samples are shown as red data points. The CGE1 bait is shown in blue, known CGE1 interaction partners in green.

We next ran LC-MS/MS analyses on streptavidin eluates obtained from three independent experiments for the four strains under non-stress and heat stress conditions and identified a total of 1169 protein groups (Supplemental Data Set S1). We first filtered for proteins that were identified in at least one strain and one condition in all three replicates. After median normalization, we next filtered for proteins significantly enriched at least two-fold in the TurboID-CGE1- and mCherry-TurboID-producing lines versus the UVM4 control. This filtering step removes naturally biotinylated proteins and proteins binding unspecifically to the streptavidin beads (Mair et al., 2019) and left 89 and 76 proteins enriched in TurboID-CGE1 versus UVM4 under non-stress and heat stress conditions, respectively (Supplemental Data Set S1). Next, we filtered for proteins that were significantly enriched in the TurboID-CGE1 versus the two mCherry-TurboID lines under each condition. This step removes proteins that get biotinylated because they are abundantly exposed to the stroma, and left ten and five proteins enriched in TurboID-CGE1 versus mCherry-TurboID under non-stress and heat stress conditions, respectively (Figure 3B). CGE1 was enriched 1997-fold under non-stress conditions and 985-fold after heat stress, whereas other significantly enriched proteins were enriched between two- and 85-fold (Supplemental Data Set S1). HSP70B and CDJ1 were significantly enriched under non-stress conditions, corroborating our immunoblot data (Figure 3A). Under heat stress, HSP70B was significantly enriched as well, while CDJ1 was significantly enriched only when compared with the line producing mCherry-TurboID at higher levels (Supplemental Data Set S1). Since we found CDJ1 enriched under both conditions in immunoblots, our filtering criteria might be somewhat too stringent. Surprisingly, we also found mitochondrial HSP70C significantly enriched under non-stress conditions.

CGE2 was significantly enriched under both conditions (Figure 3B). It received its name based on amino acid sequence motifs characteristic for chloroplast GrpE-type co-chaperones. Because of a lack of EST support and large sequence insertions in the 5’ part of the *CGE2* gene, it was not clear whether it gives rise to a functional protein (Schroda, 2004; Schroda and Vallon, 2009). Structure prediction by alpha-fold suggests a typical GrpE-fold in the C-terminal part of CGE2 but, except for a few alpha-helices, no prediction could be made for the other sequences at the N-terminus (Supplemental Figure S8A). Nevertheless, we found 29 peptides covering all parts of the sequence (Supplemental Figure S8B). Since CGE2 had the highest fold-enrichment of all proteins identified in the CGE1 proxiome and contains the typical GrpE-fold for dimerization, it is likely that CGE1 and CGE2 form heterodimers.

ACETYL-COA CARBOXYLASE 1 (ACC1), significantly enriched in TurboID-CGE1 under non-stress conditions, is a biotin-containing enzyme. Its enrichment is likely due to a more pronounced depletion of its natural biotinylation level in the mCherry-TurboID lines expressing TurboID at higher levels than the TurboID-CGE1 line (Figure 3A). The clearly cytosolic EUKARYOTIC INITIATION FACTOR 5B1 (eIF5B1) just passes the 2-fold-enrichment threshold (2.07-fold) and presumably is a false positive. Other significantly enriched proteins are a putative phytol kinase (Cre08.g381400), a putative zeta-phytoene desaturase (Cre12.g541750) and three more proteins of unknown function (Cre13.g589167, Cre15.g640250, and Cre17.g720450).

We conclude that TurboID without addition of exogenous biotin allows capturing the transient interaction of CGE1 with HSP70B in complex with co-chaperone CDJ1. It also allowed discovering CGE2 as a novel co-chaperone but it appears suitable only to a limited extent for the discovery of HSP70B substrates.

### Three labeling setups with TurboID confirm known interactions of VIPP1 with chloroplast HSP70B and interactions of VIPP1 and VIPP2 at chloroplast membranes under stress

Based on the encouraging results with TurboID-CGE1, we next wanted to identify proteins interacting with VIPP1 and VIPP2 in activities related to thylakoid biogenesis and chloroplast stress. To this end, VIPP1/2-TurboID lines, and UVM4 and mCherry-TurboID lines as controls, were grown under ambient conditions and exposed to 2 mM H_2_O_2_ for 4 h to provoke oxidative stress (Blaby et al., 2015; Theis et al., 2020). We chose three labeling protocols: first, biotin labeling *in vivo* without the addition of exogenous biotin as done for TurboID-CGE1 (Setup 1). Second, biotin labeling *in vitro*, where 500 µM biotin together with 2.5 mM ATP and an ATP-regenerating system were added to crude membrane extracts for 30 min (Setup 2). Membrane extracts were prepared from cells exposed or not to H_2_O_2_ for 4 hours. Third, biotin labeling *in vivo* with the addition of 1 mM biotin to the cultures for 4 h in parallel to H_2_O_2_ exposure (Setup 3).

We first analyzed biotinylated proteins in inputs and streptavidin eluates by immunoblotting. For Setup 1 (Figure 4A), we observed weak cis-biotinylation of the baits, reduced biotinylation of naturally biotinylated proteins correlating with the expression level of the baits, and little trans-biotinylation when compared with UVM4. Exogenously added biotin strongly enhanced cis- and trans-biotinylation (Setups 2 and 3, Figure 4A) and, in Setup 3, also appeared to enhance natural protein biotinylation.

**Figure 4.**
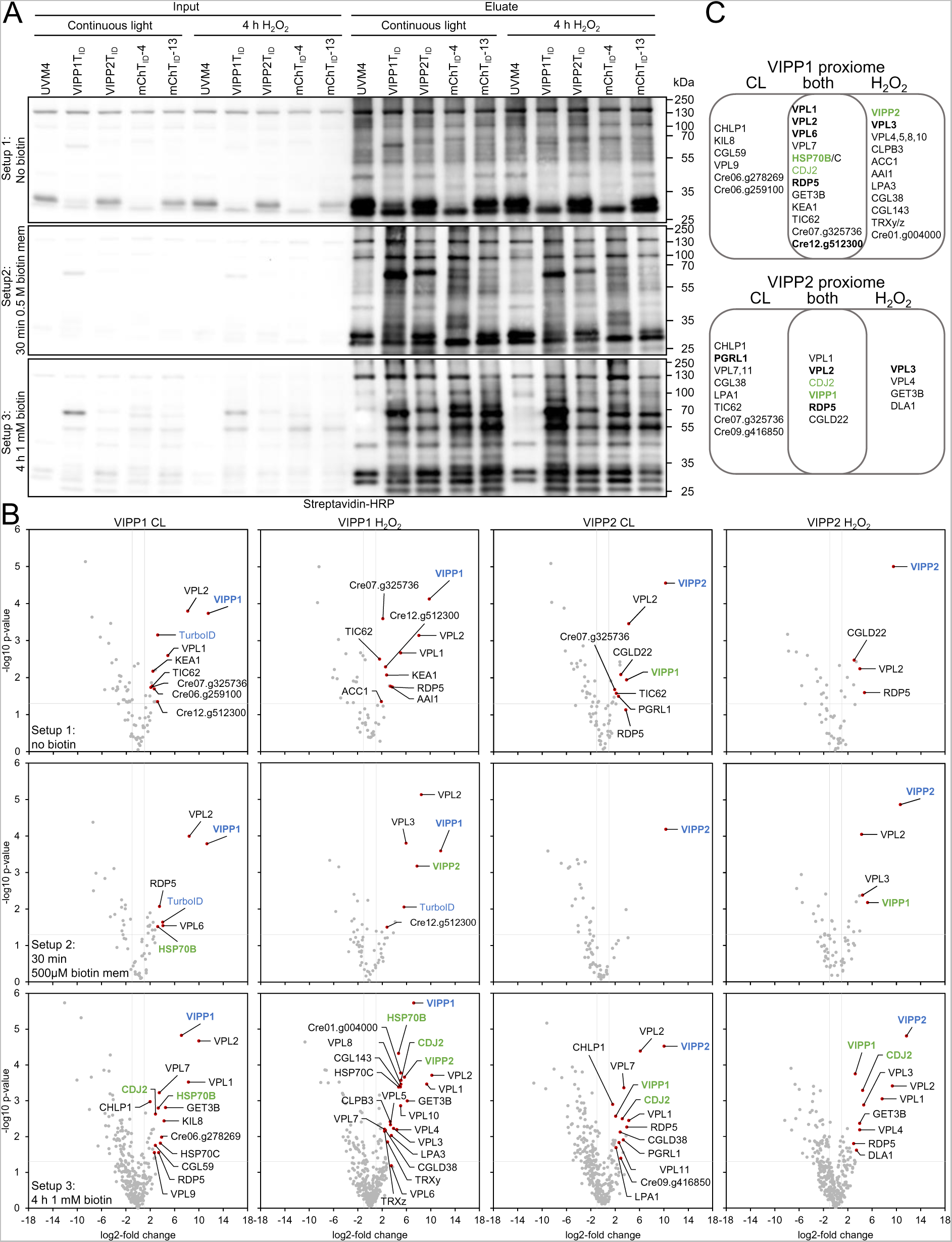
TurboID-based *in-vivo* proximity labeling using VIPP1 and VIPP2 as baits. (**A**) Immunoblot analysis. Cultures of the UVM4 recipient strain, transformants producing VIPP1TID and VIPP2T_ID_, and two transformants accumulating different levels of mCherryT_ID_ were grown to mid-log phase at 22°C (CL). Half of the cultures was exposed to 2 mM H_2_O_2_ for 4 h. Before the treatments, cells were supplemented without biotin (Setup 1) or with 1 mM biotin (Setup 3), harvested, and lysed. Alternatively, 500 µM biotin was added for 30 min to membrane fractions following the H_2_O_2_ treatment and removed again by passage through PD-10 desalting columns (Setup 2). Biotinylated proteins were affinity-purified with streptavidin beads. Protein extracts before incubation (input) and after elution from the beads (eluate) were separated by SDS-PAGE and analyzed by immunoblotting with streptavidin-HRP. One of three biological replicates each is shown. **(B)** Volcano plots of the VIPP1 and VIPP2 proxiomes resulting from Setups 1-3. Streptavidin bead eluates of the samples shown in (A) were analyzed by mass spectrometry. Shown is a comparison between protein abundances of the VIPP1/2T_ID_ samples and two T_ID_-mCherry lines after subtraction of contaminants and endogenous biotinylated proteins from the UVM4 control strain. Proteins significantly enriched in the VIPP1/2T_ID_ samples are shown as red data points. VIPP1/2 baits are shown in blue, known VIPP1/2 interaction partners in green. **(C)** Venn diagram showing proteins of the VIPP1/2 interactomes identified only under non-stress conditions (CL), only after H_2_O_2_ treatment, or under both conditions. Proteins in bold were identified in at least two of the three labeling approaches, proteins in green are known VIPP1/2 interaction partners.

LC-MS/MS analysis of streptavidin eluates performed in biological triplicates resulted in the identification of a total of 897 (Setup 1), 477 (Setup 2), and 1464 (Setup 3) protein groups (Supplemental Data Sets S2-4). These numbers perfectly meet the expectation from the immunoblot data that exogenously added biotin boosts protein biotinylation and that protein complexity is lower when focusing on membranes. Accordingly, following the same filtering steps employed for TurboID-CGE1, the largest number of proteins significantly enriched in VIPP1/2-TurboID was for Setup 3 with 10-22 proteins, followed by Setup 1 with 4-10 proteins and Setup 2 with 1-6 proteins (Figure 4B). VIPP1/2-TurboID baits were enriched between 724- and 3246-fold (Table 1; Supplemental Data Sets S2-4). Only for Setup 3, the enrichment for VIPP1-TurboID was markedly lower (140- to 148-fold). This is due to a stronger labeling of VIPP1 in the mCherry-TurboID control lines, presumably because of an enhanced labeling rate in this setup combined with the relatively high abundance of VIPP1 (0.05% of total cell proteins, which corresponds to a less abundant Calvin-Benson-Cycle enzyme (Liu et al., 2007; Hammel et al., 2020)). The enrichment for non-bait proteins ranged between 3- and 1163-fold (Table 1).

**Table 1.**
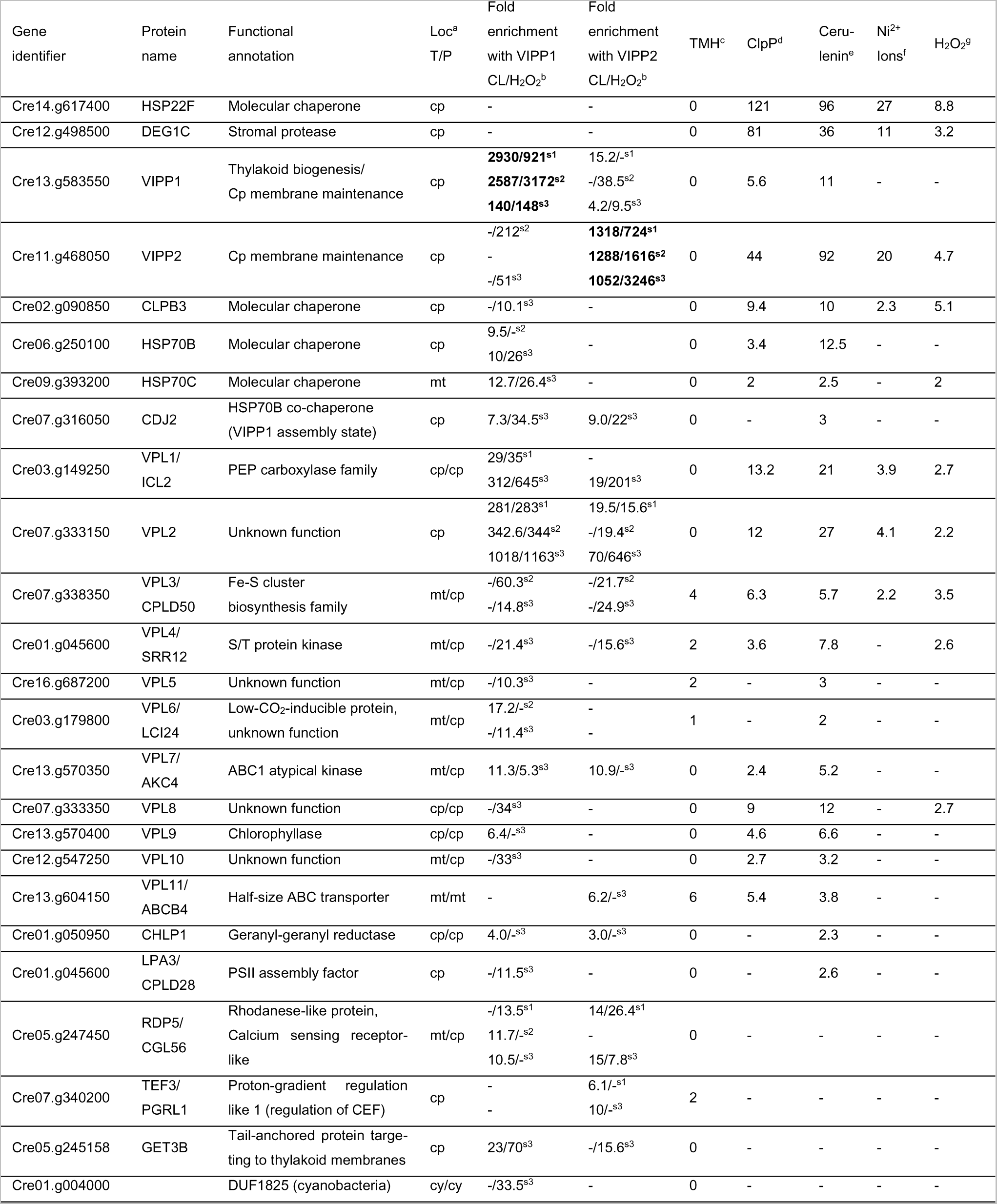
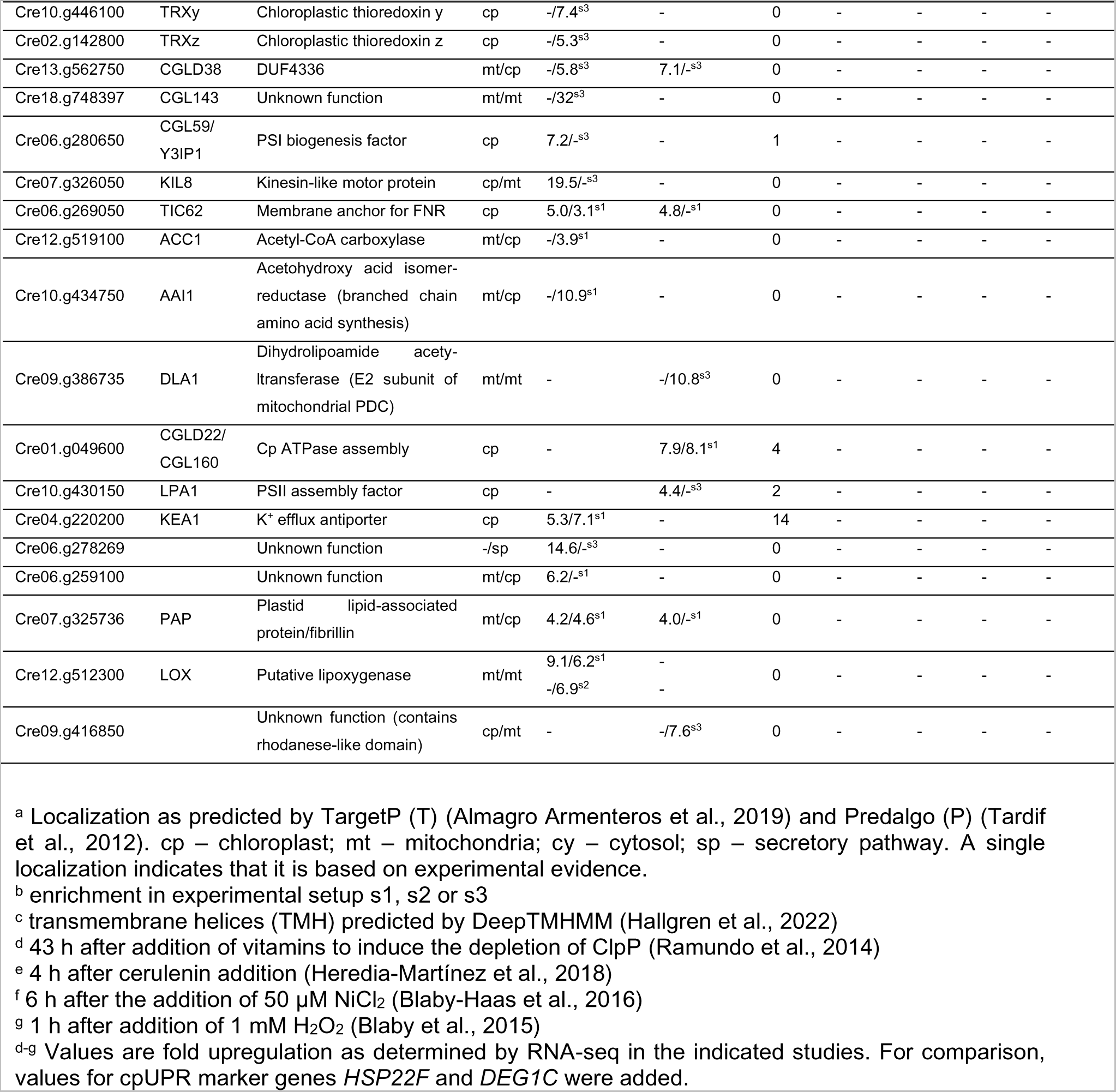
List of proteins in the VIPP1/2 proxiomes and upregulation of their encoding genes by chloroplast stresses from previous studies.

We first looked whether the LC-MS/MS data can confirm known interactions. We found VIPP2 enriched with VIPP1-TurboID under Setups 2 and 3 only after H_2_O_2_ treatment (Figures 4B and 4C; Table 1). This is expected, since VIPP2 is barely expressed under ambient conditions and strongly upregulated under stress (Nordhues et al., 2012; Ramundo et al., 2014; Perlaza et al., 2019; Theis et al., 2020). Conversely, VIPP1 was enriched with VIPP2-TurboID in all three setups under ambient and H_2_O_2_ stress conditions. The enrichment of VIPP1 with VIPP2-TurboID and VIPP2 with VIPP1-TurboID in Setup 2 suggests that both proteins form heterooligomeric complexes at membranes under H_2_O_2_ stress, corroborating previous AP-MS data (Theis et al., 2020). Other proteins known to interact with VIPP1 are HSP70B and its J-domain co-chaperone CDJ2 (Liu et al., 2007). Both were found to be enriched with VIPP1-TurboID in Setup 3 under ambient and H_2_O_2_ stress conditions. HSP70B was enriched also in Setup 2 under ambient conditions, suggesting an interaction at chloroplast membranes. Interestingly, we found CDJ2 but not HSP70B enriched with VIPP2-TurboID in Setup 3 under both conditions. In Setup 3, we also found mitochondrial HSP70C to be enriched with VIPP1-TurboID to the same extent as HSP70B under non-stress and H_2_O_2_ stress conditions.

In summary, exogenously added biotin strongly enhanced TurboID-mediated protein biotinylation and biotinylation of naturally biotinylated proteins. *In vivo* labeling and *in vitro* labeling on membrane extracts confirms a formation of VIPP1-VIPP2 heterooligomers at membranes under stress and confirms the interaction between VIPPs and the chloroplast HSP70 system.

### Genes encoding 17 proteins in the proxiomes of VIPP1 and VIPP2 were upregulated under chloroplast stress

Since we were interested in identifying VIPP1/2 interacting proteins with functions related to chloroplast stress, we wondered whether the genes encoding the 39 proteins significantly enriched with VIPP1/2-TurboID were responsive to conditions provoking chloroplast stress. To elucidate this, we consulted previous RNA-seq studies monitoring genes up-regulated after depletion of ClpP (Ramundo et al., 2014), after addition of chloroplast fatty acid synthesis inhibitor cerulenin (Heredia-Martínez et al., 2018), and after the addition of Ni^2+^ ions (Blaby-Haas et al., 2016) or H_2_O_2_ (Blaby et al., 2015). 17 of the 39 genes were upregulated under at least one of these stresses (Table 1). Among these were genes encoding (co-)chaperones HSP70B, HSP70C, CDJ2, and CLPB3. CLPB3 was enriched with VIPP1-TurboID only in Setup 3 and only under H_2_O_2_ stress. It functions as a protein disaggregase in the chloroplast and localizes to punctae situated next to the thylakoid membrane system (Kreis et al., 2022). Of the remaining 13 genes, only two encoded proteins with a clear functional annotation: CHLP1, which was enriched with VIPP1- and VIPP2-TurboID only under ambient conditions, and LPA3 (LOW PS II ACCUMULATION 3), which was enriched only with VIPP1 under H_2_O_2_ stress. CHLP1 is a geranylgeranyl reductase that catalyzes the reduction of geranylgeranyl diphosphate to phytyl diphosphate providing phytol for tocopherol and chlorophyll biosynthesis (Tanaka et al., 1999). LPA3 acts together with LPA2 in the stable assembly of CP43 into the photosystem II core complex (Chi et al., 2012). As no clear functional annotation exists for the proteins encoded by the last 11 genes we named them VIPP PROXIMITY LABELING (VPL1-11). The most highly enriched among these were VPL1-3, with maximum enrichment factors of 645-, 1163-, and 60-fold, respectively (Table 1). They showed the same enrichment pattern with VIPP1- and VIPP2-TurboID, with VPL1 and VPL2 enriched under ambient and H_2_O_2_ stress conditions, while VPL3 was enriched only under H_2_O_2_ stress (Figure 4C). Enrichment in Setup 2 of VPL2 and VPL3 suggests that they interact with the VIPPs also at membranes (VPL3 has three predicted transmembrane helices while VPL2 has none). VPL1 and VPL3 are conserved in the green lineage, while VPL2 is conserved only in Chlorophyceae (Supplemental Figure S9B). Although VPL1 was annotated as an isocitrate lyase, there is no experimental evidence for this function and ICL2 shows only 38% sequence identity with mitochondrial ICL1. No functional annotation exists for VPL2. Alpha-fold predicts VPL2 to contain 4-5 extended α-helices with the propensity to form coiled-coils, interrupted by unstructured regions (Supplemental Figure S9A). VPL3/CPLD50 is annotated as member of the Fe-S cluster biosynthesis family but experimental evidence is missing. VPL4-10 are enriched with VIPP1-TurboID, VPL4 and VPL7 also with VIPP2-TurboID and VPL11 is only enriched with VIPP2-TurboID. Among these, VPL4, 5, 8, and 10 were enriched only under H_2_O_2_ stress. According to interPro (Blum et al., 2021), VPL5, 6, 8, and 10 have no functional domains, VPL4 and VPL7 are kinases, VPL9 is a chlorophyllase, cleaving off the phytol tail from chlorophyll, and VPL11 is a half-size ABC transporter recently named ABCB4 (Li et al., 2022). Predalgo predicts VPL11/ABCB4 to localize to mitochondria, while VPL1-10 are all predicted to localize to the chloroplast.

In summary, 17 proteins in the VIPP1/2 proxiome are encoded by genes that were upregulated under chloroplast stress conditions. 11 of them lack a clear functional annotation and were named VPL1-11.

### Two large groups of proteins in the VIPP1/2 proxiomes are involved in the biogenesis of thylakoid membrane protein complexes and in the regulation of photosynthetic electron flow

22 more proteins were significantly enriched with VIPP1/2-TurboID whose encoding genes were not induced by the four chloroplast stresses. Four of them can be grouped into proteins that are involved in various assembly processes and include LOW PS II ACCUMULATION 1 (LPA1, PS II assembly, (Peng et al., 2006)), YCF3 INTERACTING PROTEIN 1 (Y3IP1, PS I assembly, (Albus et al., 2010)), CONSERVED IN GREEN LINEAGE 160 (CGL160, chloroplast ATP synthase assembly, (Fristedt et al., 2015)), and GUIDED ENTRY OF TAIL-ANCHORED PROTEINS 3 (GET3B, targeting of tail-anchored proteins to thylakoids, (Anderson et al., 2021)). Also LPA3 and CHLP1 from the chloroplast stress-inducible proteins can be included here.

A second group comprises proteins involved in the regulation of photosynthetic electron flow, including TRANSLOCON AT THE INNER ENVELOPE 62 (TIC62), PROTON GRADIENT REGULATION LIKE 1 (PGRL1), potentially RHODANESE DOMAIN PROTEIN 5 (RDP5), and two thioredoxins. TIC62 anchors ferredoxin-NADP(H) oxidoreductase (FNR) to the thylakoid membrane (Benz et al., 2009), which influences the speed at which photosynthetic control is induced and therefore plays a role in alleviating high light stress (Rodriguez-Heredia et al., 2022). PGRL1 together with PGR5 is involved in antimycin A-sensitive cyclic electron flow (DalCorso et al., 2008). RDP5 is related to the Ca^2+^-sensing receptor (CAS) protein and, although it has no predicted transmembrane domain, was significantly enriched with VIPP1-TurboID at membranes (Figure 4B, Setup 2). Thylakoid-localized CAS relays a Ca^2+^ signal to a retrograde signal essential for maintaining the expression of genes important for operating the CO_2_-concentrating mechanism that provides a sink for electrons from the light reactions (Wang et al., 2016). CAS also links a Ca^2+^ signal to the high light-induced expression of the *LHCSR3* gene that is crucial for nonphotochemical quenching (Petroutsos et al., 2011). Thioredoxin z (TRXz) has been shown to activate Calvin-Benson cycle protein phosphoribulokinase *in vitro* (Le Moigne et al., 2021). Plant TRXy interacts with 2-Cys peroxiredoxins but the physiological relevance of this interaction is not clear (Jurado-Flores et al., 2020).

Several significantly enriched proteins cannot be functionally grouped, including K^+^-EFFLUX ANTIPORTER 1 (KEA1), which is located to the chloroplast envelope and together with KEA2 plays a critical role for the rapid downregulation of stromal pH, especially during light–dark transitions (Aranda Sicilia et al., 2021); ACETOHYDROXY ACID ISOMEROREDUCTASE (AAI1), which catalyzes the second step in branched chain amino acid synthesis (Vallon and Spalding, 2009); DIHYDROLIPOAMIDE ACETYLTRANSFERASE (DLA1), which is the E2 subunit of mitochondrial pyruvate decarboxylase (Bohne et al., 2013); a kinesin-like motor protein (KIL8); a putative plastid lipid-associated protein (PAP); and a putative lipid peroxidase (LOX).

Six proteins significantly enriched with VIPP1/2-TurboID have no clear functional annotation (CGLD38, CGL143, Cre01.g004000, Cre06.g278269, Cre06.g259100, and Cre09.g416850). Biotin-containing enzyme ACC1 was (mildly) enriched with VIPP1-TurboID, presumably due to a more pronounced depletion of its endogenous biotinylation level in the mCherry-TurboID lines, as postulated for the TurboID-CGE1 experiment.

Among the 39 proteins significantly enriched with VIPP1/2-TurboID, 32 were predicted by TargetP (Almagro Armenteros et al., 2019) and/or Predalgo (Tardif et al., 2012) to be targeted to the chloroplast, five were predicted by both programs to be targeted to mitochondria (HSP70C, VPL11/ABCB3, CGL143, DLA1, LOX), and one each to be targeted to cytosol (Cre01.g004000) and secretory pathway (Cre06.g278269) (Table 1).

In summary, in addition to the group of proteins associated with chloroplast stress, other proteins in the VIPP1/2 TurboID proxiomes are involved in the biogenesis of thylakoid membrane protein complexes and the control of photosynthetic electron flow and have individual functions.

### Reciprocal VPL2-TurboID confirms the proximity of VPL2 to VIPP1

To exemplarily confirm the results from VIPP1/2-TurboID PL, we selected VPL2 as a bait for a reciprocal labeling experiment. VPL2 was chosen because it was one of the most enriched proteins with VIPP1-TurboID under all three experimental setups (Table 1), its predicted amino acid sequence was well supported with identified tryptic peptides (Supplemental Figure S9B), and it has no predicted transmembrane domains. We synthesized the VPL2 coding sequence, interrupted by *RBCS2* introns 1 and 2 and including the predicted transit peptide, with optimized codon usage and assembled it into a level 0 construct for MoClo (Supplemental Figure S1). Following the design used for APEX2 constructs, we assembled level 2 constructs for the production of VPL2-TurboID (Figure 5A). We identified several transformants accumulating VPL2-TurboID in a single protein band and chose one with VPL2-TurboID expression levels between those of the two mCherry-TurboID control lines (Supplemental Figure S10).

**Figure 5.**
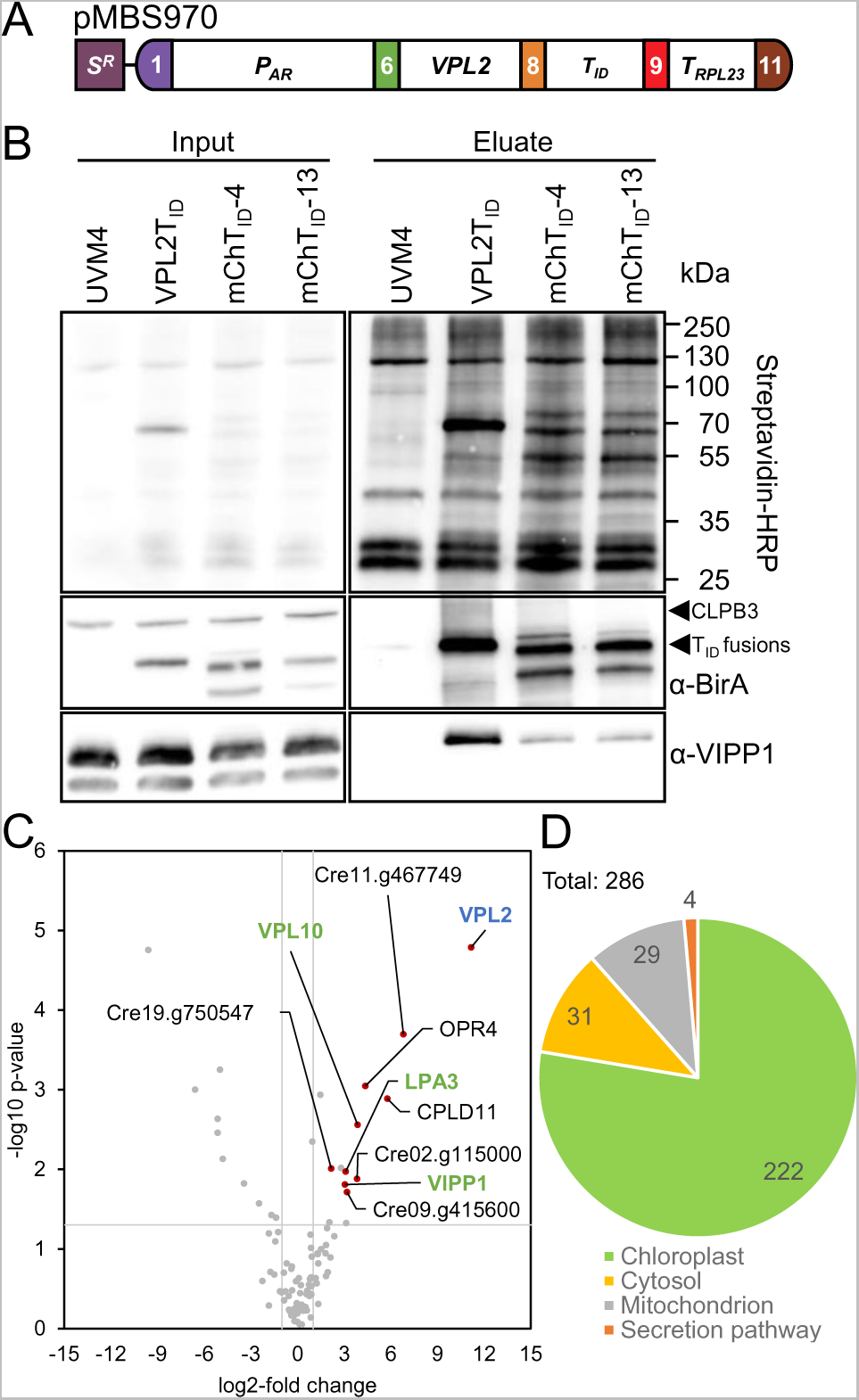
TurboID-based *in-vivo* proximity labeling using VPL2 as a bait. (**A**) Level 2 construct conferring resistance to spectinomycin (*S^R^*) and driving the production of VPL2 with a C-terminal TurboID fusion as described in Figure 1A. (**B**) Immunoblot analysis. Cultures of the UVM4 recipient strain, a transformant producing VPL2T_ID_, and two transformants accumulating different levels of mCherryT_ID_ were grown to mid-log phase at 22°C. Cells were supplemented with 1 mM biotin for 4 h, harvested, and lysed. Biotinylated proteins were affinity-purified with streptavidin beads. Protein extracts before incubation with beads (input) and after elution from the beads (eluate) were separated by SDS-PAGE and analyzed by immunoblotting with streptavidin-HRP and antibodies against BirA and VIPP1. One of three biological replicates is shown. **(C)** Volcano plot of the VPL2 proxiome. Streptavidin bead eluates of the samples shown in (B) were analyzed by mass spectrometry. Shown is a comparison between protein abundances of the VPL2T_ID_ sample and two T_ID_-mCherry lines after subtraction of contaminants and endogenously biotinylated proteins from the UVM4 control strain. Proteins significantly enriched in the VPL2T_ID_ sample are shown as red data points. The VPL2 bait is shown in blue, proteins found before in the VIPP1/2 proxiomes in green. (**D**) Pie diagram showing the predicted localizations of all 286 proteins found to be significantly enriched in lines producing TurboID fusions with CGE1, VIPP1/2, and VPL2 against UVM4.

Biotin labeling was done according to experimental Setup 3 (addition of 1 mM biotin for 4 h to cultures with the UVM4 control, two mCherry-TurboID lines, and the VPL2-TurboID line). Since VPL2 was not further enriched with VIPP1-TurboID under H_2_O_2_ stress (Table 1), we only went for ambient conditions. Immunoblot analysis with streptavidin-HRP revealed strong cis-biotinylation of VPL2-TurboID and successful enrichment of VPL2-TurboID, mCherry-TurboID, and trans-biotinylated proteins with the streptavidin beads (Figure 5B). Again, the biotin boost rescued reduced biotinylation of naturally biotinylated proteins. Immunoblot analysis with the VIPP1 antibody revealed strong enrichment of VIPP1 with VPL2-TurboID. Less VIPP1 was enriched with mCherry-TurboID and none when TurboID was absent (UVM4 control).

LC-MS/MS analysis on streptavidin eluates obtained from three independent experiments for the four strains resulted in the identification of a total of 831 protein groups (Supplemental Data Set S5). Following the same filtering steps employed for TurboID-CGE1, we found a 2298-fold enrichment of VPL2 and an enrichment for non-bait proteins ranging between 4.5- and 112-fold (Table 2). Nine proteins were significantly enriched with VPL2-TurboID (Figure 5C) and all were predicted to be targeted to the chloroplast by TargetP and/or Predalgo (Table 2). Among them were VIPP1, VPL10, and LPA3, which had been found to be significantly enriched with VIPP1/2-TurboID. Genes encoding these three proteins and that of a putative glucan 1,4-alpha-glucosidase were all upregulated under at least one condition provoking chloroplast stress (Table 2). Other significantly enriched proteins include NADH:plastoquinone reductase NDA2, which is involved in the antimycin A insensitive pathway for cyclic electron flow and in chlororespiration (Jans et al., 2008); CGLD11, which is involved in F_1_ assembly during the biogenesis of chloroplast ATP synthase in Arabidopsis (Grahl et al., 2016) (this protein must be handled with care as its identification and quantification is based only on a single peptide); octatricopeptide repeat protein (OPR4); and two more proteins of unknown function (Table 2).

**Table 2.** List of proteins in the VPL2 proxiome and upregulation of their encoding genes by chloroplast stresses from previous studies.

| Gene identifier | Protein name | Functional annotation | Loc<sup>a</sup><br>T/P | Fold enrichment with VPL2<sup>b</sup> | TMH<sup>c</sup> | ClpP<sup>d</sup> | Cerulenin<sup>e</sup> | Ni<sup>2+</sup> ions<sup>f</sup> | H<sub>2</sub>O<sub>2</sub><sup>g</sup> |
| --- | --- | --- | --- | --- | --- | --- | --- | --- | --- |
| Cre07.g333150 | VPL2 | Unknown function | cp | **2298** | 0 | 12 | 27 | 4.1 | 2.2 |
| Cre13.g583550 | VIPP1 | Thylakoid biogenesis/<br>Cp membrane maintenance | cp | 8.3 | 0 | 5.6 | 11 | - | - |
| Cre12.g547250 | VPL10 | Unknown function | mt/cp | 14.6 | 0 | 2.7 | 3.2 | - | - |
| Cre01.g045600 | LPA3/<br>CPLD28 | PSII assembly factor | cp | 8.6 | 0 | - | 2.6 | - | - |
| Cre09.g415600 |  | Glucan<br>1,4-alpha-glucosidase | cp/cp | 9.1 | 0 | 2.6 | - | - | - |
| Cre02.g106700 | OPR4 | Octotricopeptide repeat<br>protein with RAP domain | cp/mt | 20.7 | 0 | - | - | - | - |
| Cre02.g115000 |  | Unknown function | cp/cp | 14.2 | 0 | - | - | - | - |
| Cre11.g467749 |  | Unknown function | cp/cp | 112 | 0 | - | - | - | - |
| Cre19.g750547 | NDA2 | Type-II NADH-plastoquinone reductase | mt/cp | 4.5 | 0 | - | - | - | - |
| Cre08.g372000 | CGLD11/<br>BAF3 | Cp ATPase CF1 assembly | cp/cp | 54.8<br>(only 1 peptide) | 0 | - | - | - | - |

<sup>a</sup> Localization as predicted by TargetP (T) (Almagro Armenteros et al., 2019) and Predalco (P) (Tardif et al., 2012). cp – chloroplast; mt – mitochondria. A single localization indicates that it is based on experimental evidence.
<sup>b</sup> enrichment in experimental Setup 3
<sup>c</sup> transmembrane helices (TMH) predicted by DeepTMHMM (Hallgren et al., 2022)
<sup>d</sup> 43 h after addition of vitamins to induce the depletion of ClpP (Ramundo et al., 2014)
<sup>e</sup> 4 h after cerulenin addition (Heredia-Martínez et al., 2018)
<sup>f</sup> 6 h after the addition of 50 μM NiCl<sub>2</sub> (Blaby-Haas et al., 2016)
<sup>g</sup> 1 h after addition of 1 mM H<sub>2</sub>O<sub>2</sub> (Blaby et al., 2015)
<sup>d-g</sup> Values are fold upregulation as determined by RNA-seq in the indicated studies.

We conclude that reciprocal TurboID with VPL2, present in the VIPP1/2 proxiomes, resulted in significant enrichment of VIPP1. VPL10 and LPA3 from the VIPP1/2 proxiomes were also enriched, further confirming the VIPP1/2 TurboID data.

### Using PL data to decipher the chloroplast proteome

PL not only allows identifying the proxiome of a particular bait, but also provides information on the composition of a compartment’s proteome (Rhee et al., 2013; Kim et al., 2014; Mair et al., 2019). To get this information, we combined all proteins that were significantly enriched in the streptavidin eluates on extracts of transformants expressing our TurboID-tagged baits against the eluates from the UVM4 control strain. Here we filter out contaminants and natively biotinylated proteins and should retain only proteins that were biotinylated by TurboID activity in the chloroplast. Given the incomplete chloroplast-targeting of mCherry-TurboID, as judged from the accumulation of a protein band presumably derived from the precursor protein, the mCherry data were omitted and only those with CGE1, VIPP1, VIPP2, and VPL2 as baits considered. This resulted in a total of 286 proteins. Of these, 222 (78%) were predicted to localize to the chloroplast based mainly on Predalgo predictions (Tardif et al., 2012) and manual curation (Westrich et al., 2021). 31 proteins were predicted to be mitochondrial, 29 to be cytosolic, and four to be secreted (Figure 5D).

## Discussion

### APEX2 and BioID are not suitable for PL in the *Chlamydomonas* chloroplast

Here we report on the development of proximity labeling (PL) for studying protein interaction networks in the *Chlamydomonas* chloroplast. We have applied the three most commonly used PL systems based on APEX2 (Lam et al., 2015), BioID (Roux et al., 2012), and TurboID (Branon et al., 2018) and a total of five stromal and membrane-peripheral baits (CGE1, VIPP1, VIPP2, VPL2, and mCherry as control). APEX2 fused to CGE1 did not result in protein biotinylation after biotin-phenol (BP) addition to *Chlamydomonas* cell cultures even if longer incubation times with BP or H_2_O_2_ were employed (Figure 1B; Supplemental Figure S3). Presumably, *Chlamydomonas* cells are not permeable for BP. The failure of APEX2-mediated PL was also reported in a parallel submission establishing PL in *Chlamydomonas* (Lau et al., 2022). Cell permeability problems have been reported also in some mammalian cell types and tissues, in thicker *Drosophila* tissues, or in fission yeast. While labeling could eventually be achieved in mammalian cells by increasing BP concentrations from 0.5 mM to 2.5 mM and higher (Tan et al., 2020), this was not sufficient in fission yeast, where cell permeabilization by increased osmolarity was required (Hwang and Espenshade, 2016). In *Drosophila*, small amounts of detergent were required to increase permeability (Mannix et al., 2019). Since PL in chloroplasts requires BP to traverse three membranes, we did not pursue APEX2-mediated PL further, albeit we could demonstrate rapid labeling in soluble cell extracts (Figure 1C).

Cis-biotinylation of BioID fused to CGE1 was only clearly detectable after 24 hours of labeling time and was more pronounced when 1 mM biotin was used compared with 500 µM (Figure 2C). Compared to TurboID fusions, trans-biotinylation was hardly detectable (Figure 2C). Long labeling times of 22-48 h and biotin concentrations ranging between 50 µM and 2 mM also were required in studies employing BioID fusions in cytoplasm/nucleus or at plasma membranes of *N. benthamiana*, tomato, rice, and Arabidopsis cells but in these studies BioID-mediated protein biotinylation was much higher than in *Chlamydomonas* (Lin et al., 2017; Conlan et al., 2018; Khan et al., 2018; Das et al., 2019; Mair et al., 2019; Zhang et al., 2019; Arora et al., 2020). This suggest that BioID is at most of limited use for PL in the chloroplast of *Chlamydomonas*.

### Differences and commonalities of TurboID-mediated PL in *Chlamydomonas* and land plants

In contrast to APEX2 and BioID, TurboID fused to any of the five baits resulted in efficient protein biotinylation in the *Chlamydomonas* chloroplast. We observed clearly detectable cis- and trans-biotinylation even in the absence of exogenous biotin (Figures 2B, 2C, 3A, 4A; Supplemental Figure S7B). However, the addition of biotin to the cultures strongly enhanced protein biotinylation even under H_2_O_2_ stress conditions, and we found biotin concentrations of 1 mM and labeling times of 1-6 h to be optimal (Figures 2C, 4A). Background biotinylation in the absence of biotin was observed also for TurboID fusions with three Calvin-Benson-Cycle enzymes in *Chlamydomonas* in a parallel study (Lau et al., 2022). In that study, similar labeling times and biotin concentrations as used by us were found to be optimal even in another strain background, demonstrating the reproducibility of TurboID-mediated PL in the chloroplast of *Chlamydomonas*.

TurboID applications to identify protein-protein interaction networks in Arabidopsis and *N. benthamiana* used only 50-200 µM of exogenously added biotin but similar labeling times of 0.5-12 h, although strong protein biotinylation was observed already after as little as 15 min (Mair et al., 2019; Zhang et al., 2019; Xu et al., 2021; Tang et al., 2022; Wurzinger et al., 2022). While in these studies no effects on naturally biotinylated proteins was reported, we observed strong effects in *Chlamydomonas* with loss of natural protein biotinylation correlating with TurboID expression levels (Figures 2B, 2C, 3A, 4A; Supplemental Figure S7B). Hence, the high activity of TurboID effectively competes with natural biotinylation in the chloroplast. Unexpectedly, we observed no negative effects on growth and no increased accumulation of cpUPR marker proteins in TurboID-expressing lines compared to the wild type even if cells were subjected to 24 h heat stress or 10 h of high light (Supplemental Figure 7). Nevertheless, the reduced natural biotinylation should be kept in mind when using TurboID. This is also because a greater reduction in natural biotinylation in control cells compared with bait cells can lead to an enrichment of naturally biotinylated proteins in bait cells, as was observed with ACC1 (Figures 3B, 4B). Importantly, natural biotinylation could be rescued to some extent by the addition of 1 mM exogenous biotin for up to 6 hours, which also abolished ACC1 enrichment (Figures 2C, 4A, 5B). The expression of TurboID-baits should not be too high to avoid too much interference with natural biotinylation. We estimate the expression level achieved in fusions of TurboID with CGE1 and VIPP1 to be 0.01-0.05% of total cell proteins, based on the accumulation of the proteins to levels similar to the native proteins (Liu et al., 2007). However, it should be noticed that effective biotinylation was observed with VIPP2-TurboID, which was expressed at much lower levels than TurboID-CGE1 and VIPP1-TurboID (Figure 2B).

In land plants, a desalting step was essential to remove excess biotin from protein extracts prior to incubation with streptavidin beads (Mair et al., 2019; Zhang et al., 2019; Arora et al., 2020; Xu et al., 2021; Tang et al., 2022; Wurzinger et al., 2022). This desalting step was required for *Chlamydomonas* only if biotin was added to crude membrane extracts. If added to the cell culture, the routine cell harvesting protocol including one washing step removes excess biotin sufficiently.

### Boosting protein biotinylation by adding biotin improves the power of TurboID to reveal protein interaction networks

We performed TurboID-mediated PL and mass spectrometry analysis with three experimental setups: in Setup 1 no biotin was added to the cultures; in Setup 2, biotin was added to crude membrane extracts; and in Setup 3, biotin was added to the cultures. All three setups were used for VIPP1/2-TurboID fusions and therefore allow comparisons to be made (Figure 4, Supplemental Data Sets S2-S4). The number of identified biotinylated proteins was highest in Setup 3 (1464 proteins), followed by Setup 1 (897 proteins) and Setup 2 (477 proteins). The 1.6-fold increase in the number of proteins identified in Setup 3 compared with Setup 1 suggests that the addition of biotin to the cultures greatly increases TurboID-mediated specific and background biotinylation. The number of proteins identified could be even higher if a mass spectrometer with an ion trap (C-trap, trapped ion mobility spectrometry) had been used; we used a TripleTOF instrument without an ion trap. In a first filtering step comparing the proteins identified in the TurboID lines with the wild type, about 90% of the proteins were removed. These are naturally biotinylated proteins and contaminants that bind to the streptavidin beads. In a second filtering step comparing the proteins identified in the bait lines with the mCherry controls, another about 90% of the proteins were removed. These are proteins that get biotinylated because they are abundantly present in the same compartment as the bait and/or expose readily accessible primary amines (Mair et al., 2019; Zhang et al., 2019; Mair and Bergmann, 2021). As pointed out by Mair et al. (2019), the non-bait-TurboID control should have approximately the same expression level as the bait-TurboID fusion. We used two controls expressing mCherry-TurboID at different levels. This might be too stringent, as specific interaction partner CDJ1 in the CGE1 proxiome was enriched when using one control but not with two (Supplemental Data Set S1).

Even after the two filtering steps, the number of significantly enriched proteins in Setup 3 was 1.6- to 2.5-fold higher than in Setup 1. Thus, the larger number of biotinylated proteins in Setup 3 compared to Setup 1 also resulted in a proportionally larger number of significantly enriched proteins. More candidates improve statistical power, as shown by the enrichment of TurboID itself in Setup 1 and 2, but not in Setup 3 (Figure 4B). This is consistent with previous results where most FAMA transcription factor interaction candidates were identified by TurboID-mediated PL after 3 hours of labeling time compared with 0.5 hours of labeling time (Mair et al., 2019). Nevertheless, experimental Setup 1 allowed the identification of known CGE1 partner proteins HSP70B and CDJ1 and of the obvious new partner CGE2 (Figure 3B). Setup 1 also allowed the identification of the highly enriched proteins VPL1, VPL2, and RDP5 in the VIPP1/2 proxiomes (Figure 4; Table 1). However, Setup 1 only allowed probing of the known interaction between VIPP1 and VIPP2, but not the known interaction of VIPP1 with HSP70B and CDJ2, which were probed in Setup 3 (Figure 4). Hence, consistent with the findings of Mair et al. (2019), Setup 3 appears to give a more comprehensive picture on a protein interaction network than Setup 1.

The strong enrichment of biotinylated VIPP1/2-TurboID and even mCherry-TurboID in Setup 2 suggests that at least cis-biotinylation could be enhanced in crude membrane extracts (Figure 4A). However, because no proteins were enriched in Setup 2 in addition to those found in Setups 1 and 3, this *in vitro* PL setup has no additional benefit for proxiome mapping compared with the *in vivo* PL setups. It is also problematic here that mCherry-TurboID is no good control because it is present in membrane fractions only as a contaminant from soluble proteins, whereas VIPP1/2-TurboID are truly membrane-associated.

### Not all interactions are probed by TurboID-mediated PL

Despite the ability of PL in Setup 3 to probe known interactions of VIPP1 with VIPP2, HSP70B, and CDJ2, other known interactions of VIPP1 with CGE1, HSP90C, and HSP22E/F (Liu et al., 2005; Heide et al., 2009; Theis et al., 2020) were not probed. In the case of CGE1, its residence time on HSP70B together with the substrate VIPP1 might be too short: GrpE-type nucleotide exchange factors bind their HSP70 partners only in the ADP state to allow rapid exchange of ADP for ATP, which triggers substrate release (Rosenzweig et al., 2019). In all *in vivo* PL datasets, HSP90C was highly enriched with all baits (including mCherry and VPL2) compared with the UVM4 control, precluding its specific enrichment with a particular bait (Supplemental Data Sets S1, S2, S4, S5). This suggests that HSP90C is readily trans-biotinylated because it is ubiquitously present in the stroma with well accessible primary amines. Alternatively, TurboID could be an HSP90C client. HSP22E accumulates at high levels under heat and oxidative stress (Rütgers et al., 2017; Theis et al., 2020), but was slightly enriched only in Setup 3 compared to the UVM4 control. Since HSP22E/F was precipitated with streptavidin beads in the heat-stressed UVM4 control (Figure 3A), it was likely removed in the first filtering step for contaminants. In addition, a large fraction of HSP22E/F may have been removed in aggregates during the precleaning centrifugation that was performed before the cell lysates were applied to the streptavidin beads. Also, in the parallel study by Lau et al. (2022), not all expected pyrenoid proteins were identified by TurboID-mediated PL. And in previous TurboID applications in land plants, not all known interactors of the bait proteins used were found either (Zhang et al., 2019; Arora et al., 2020). Possible reasons included masking of primary amines by steric hindrance or conformation, distance between bait and interactors beyond the labeling radius of activated biotin, and discrepancies between the time windows of labeling and specific interactions taking place.

These limitations of PL should also be kept in mind when PL is used for the mapping of compartment-specific proteomes, i.e., deduced from the enrichment of proteins with TurboID compared to the wild-type control (Figure 5D). There will be a bias against proteins of low abundance, proteins that lack or have a low accessibility of primary amines such as proteins deeply buried in protein complexes or in membranes, and proteins in confined subcompartments (Qin et al., 2021).

### Can the degree of enrichment be used as a measure of interaction?

The degree of enrichment of a protein in a PL experiment results from the ratio of its biotinylation in bait-TurboID cells (numerator) to mCherry-TurboID cells (denominator). The extent to which a protein is biotinylated depends on its abundance, the accessibility of primary amines on its surface, its localization, and its proximity to TurboID. An abundant protein with accessible primary amines distributed throughout the target compartment is likely to be biotinylated in mCherry-TurboID cells, contributing to a high denominator value. Even if it is enhanced biotinylated in bait-TurboID cells due to its proximity to the bait, the degree of enrichment could be moderate. In contrast, a low abundant protein with low accessibility of primary amines may not be biotinylated at all in mCherry-TurboID cells. Here, Perseus imputes a very low value for the missing value leading to a very low denominator value. Thus, even if this protein is biotinylated as much as the abundant one due to its proximity to the bait (same numerator value), it could have a much higher enrichment level. Thus, high enrichment may be due to a protein’s proximity to a bait, but it may also be due to its low abundance or low accessibility of primary amines.

Proximity to TurboID can result from interaction with the bait, but also from mere co-localization in a limited subcompartment, such as phase-separated condensates. If a protein is exclusively localized in a limited subcompartment, it may not be biotinylated in mCherry-TurboID cells, and few biotinylations by activated biotin generated by TurboID in this subcompartment may result in a very high enrichment value. In this case, strong enrichment can be achieved without close spatial proximity.

### What do we learn from the obtained proxiomes?

With TurboID we were able to probe many known interactions, including those of CGE1 with HSP70B and CDJ1 (Figure 3B) (Willmund et al., 2008), and those of VIPP1 with VIPP2, HSP70B, and CDJ2 (Figure 4B) (Liu et al., 2005; Theis et al., 2020). PL confirms the interaction of VIPP1 with VIPP2 at chloroplast membranes under oxidative stress shown previously by AP-MS (Theis et al., 2020). CLPB3 in the VIPP1 proxiome under oxidative stress conditions supports the idea that CLPB3 may aid in the removal of protein aggregates from thylakoid membranes, as proposed previously based on localization data (Kreis et al., 2022). CDJ2 in the VIPP2 proxiome suggests that the assembly state of VIPP2, like that of VIPP1, is controlled by the HSP70B/CDJ2/CGE1 chaperone system (Liu et al., 2007) (notice that VIPP2 forms rods like VIPP1 (Theis et al., 2020)). CGE2, found in the CGE1 proxiome (Figure 3B), very likely is a novel co-chaperone of CGE1, as it contains a GrpE fold for dimerization (Supplemental Figure S8), signature sequences of chloroplast GrpEs (Schroda et al., 2001), and is predicted to be targeted to the chloroplast (Schroda and Vallon, 2009). VPL2 in the proxiome of VIPP1 could be confirmed in a reciprocal PL experiment, and LPA3 and VPL10 were in the proxiomes of both, VIPP1/2 and VPL2 (Figures 4B and 5C).

Surprisingly, we found mitochondrial HSP70C in the proxiomes of CGE1 and VIPP1 (Figures 3B and 4B). In fact, HSP70C was previously found to co-precipitate with VIPP1 (Liu et al., 2005). This interaction was considered to be nonspecific and a consequence of the mixing of compartment contents during cell lysis before immunoprecipitation. Given the presence of HSP70C in the CGE1 and VIPP1 proxiomes, we must consider that HSP70C may be dually targeted to mitochondria and chloroplasts. A recent report on the localization of more than 1000 candidate chloroplast proteins by fluorescent tagging has shown that dual targeting is a common phenomenon in *Chlamydomonas* (Wang et al., 2022). Dual targeting might therefore explain why in our PL-based chloroplast proteome ∼20% of the proteins are predicted to localize to mitochondria or cytosol (Figure 5D).

In addition to probing known or expected protein interaction networks, PL has revealed several new candidate proteins with the potential to provide new insights into the function of VIPPs in the chloroplast. Such proteins have been difficult to find by conventional AP-MS based methods (Jouhet and Gray, 2009; Lo and Theg, 2012; Bryan et al., 2014). One group in the VIPP1/2 proxiomes comprises 13 proteins whose genes have been found previously to be upregulated under chloroplast stress conditions. Eleven of them lack a clear functional annotation and we named them VPL1-11. We speculate that these proteins may play a role in coping with chloroplast membrane stress and mediating retrograde signaling for the cpUPR. A second group of proteins has reported roles in the biogenesis of PS II (LPA1, LPA3), PSI (Y3IP1), and ATP synthase (CGL160, CGLD11), the targeting of tail-anchored proteins (GET3B), and the synthesis of phytol (CHLP1). A role of VIPP1 in supporting these biogenesis processes would account for the reduced levels of major thylakoid membrane protein complexes in Arabidopsis, *Chlamydomonas*, and cyanobacterial *vipp1* knockdown mutants (Kroll et al., 2001; Fuhrmann et al., 2009; Nordhues et al., 2012; Zhang et al., 2014; Zhang et al., 2016a). A third group of proteins plays roles in photosynthetic electron flow, including PGRL1, NAD2, TIC62, TRXy, TRXz, and potentially RDP5. Impaired functioning of these processes could account for the deregulation of high light-induced *LHCSR3* gene expression in *Chlamydomonas vipp1* knockdown and *vipp2* knockout lines (Nordhues et al., 2012; Theis et al., 2020). A last group contains several proteins with particular functions like KEA1.

While the involvement of VIPP1 in so many functions would explain the pleiotropic phenotypes observed in *vipp1* mutants, how can VIPP1 be involved in so many functions? We previously proposed the idea that VIPP1 might be able to organize domains in chloroplast membranes that resemble eisosomes found in fungal plasma membranes (Rütgers and Schroda, 2013; Theis and Schroda, 2016; Theis et al., 2019a; Theis et al., 2020). Local membrane bending and enrichment of specific lipid species at such domains may be required for the optimal functioning of various processes taking place at membranes (Foderaro et al., 2017).

In conclusion, TurboID-mediated PL has enabled the probing of known and new protein interaction networks in the nucleus, cytoplasm and at the plasma membrane of land plants with amazingly high sensitivity and specificity (Mair et al., 2019; Zhang et al., 2019; Arora et al., 2020; Xu et al., 2021; Tang et al., 2022). Our work and that of two parallel studies by Lau et al. (2022) and Wurzinger et al. (2022) add *Chlamydomonas* as another plant model and the chloroplast as another compartment amenable to the great power of TurboID-mediated PL. The availability of TurboID as a standard part in the *Chlamydomonas* MoClo tool kit, allowing its assembly with any bait in a single cloning step, will greatly facilitate PL in the community.

## Methods

### Strains and Culture Conditions

*Chlamydomonas reinhardtii* UVM4 cells (Neupert et al., 2009) were grown in Tris-Acetate-Phosphate (TAP) medium (Kropat et al., 2011) on a rotatory shaker at a constant light intensity of ∼40 μmol photons m^−2^ s^−1^ provided by MASTER LEDtube HF 1200 mm UO 16W830 T8 and 16W840 T8 (Philips). For heat stress experiments, exponentially growing cells were harvested by centrifugation at 3,500 g for 2 min at 25°C, resuspended in TAP medium prewarmed to 40°C, and incubated in a 40°C water bath under agitation and constant illumination at ∼40 μmol photons m^−2^ s^−1^ for 1 h. For H_2_O_2_ treatments, exponentially growing cells were incubated with 2 mM H_2_O_2_ (Sigma-Aldrich) for 4 h. Transformation was performed with the glass beads method (Kindle, 1990) as described previously (Hammel et al., 2020), with constructs linearized by EcoRV. Transformants were selected on TAP medium containing 100 µg mL^-1^ spectinomycin. Cell densities were determined using a Z2 Coulter Counter (Beckman Coulter) or photometrically by optical density measurements at 750 nm (OD750).

### Cloning of coding sequences for baits, APEX2, BioID, and TurboID

*Bait genes for level 0* – The *CGE1* gene containing all seven introns and eight exons (with exon 1 lacking sequences encoding the chloroplast transit peptide) was amplified by PCR from genomic DNA. Genomic DNA of *Chlamydomonas* strain CC-4533 was extracted as described previously (Theis et al., 2020). Four fragments were amplified to remove endogenous BsaI and BbsI restriction sites by silent mutations (primers used and product sizes are listed in Supplemental Table S1). Each fragment was flanked with BbsI restriction sites, generating unique overhangs upon BbsI digestion such that they could be directionally assembled into the pAGM1287 vector (Weber et al., 2011) during the restriction-ligation reaction (5 h at 37°C, 5 min at 50°C and 10 min at 80°C), yielding pMBS589. The *VIPP2* gene, including all nine introns and the sequences encoding the chloroplast transit peptide, was previously assembled in the same way (Theis et al., 2020). The *VIPP1* gene, including all nine introns and the sequences encoding the chloroplast transit peptide, was synthesized as described previously (Gupta et al., 2021). The 354-amino acids VPL2 protein (Cre07.g333150), including its chloroplast transit peptide, was reverse translated using the most-preferred *Chlamydomonas* codons. To enhance gene expression (Baier et al., 2018; Schroda, 2019), the first two *Chlamydomonas RBCS2* introns were inserted with the flanking sites AG/intron/GC. The sequence was split into four fragments flanked by BbsI recognition sites giving rise to distinct overhangs, synthesized (Integrated DNA Technologies), and assembled into the pAGM1287 vector in a restriction-ligation reaction, yielding pMBS969. A vector with the coding sequence for mCherry, containing the first *RBCS2* intron, was produced previously (pCM0-067) (Crozet et al., 2018). All constructs represent level 0 parts for the B3/4 position according to the Modular Cloning (MoClo) syntax for plant genes (Weber et al., 2011; Patron et al., 2015).

*APEX2, BioID, and TurboID for level 0* – The APEX2 amino acid sequence encoded by vector pcDNA3 APEX2-NES (Rhee et al., 2013; Lam et al., 2015), including a GS-linker and a FLAG-tag (DYKDDDDK) at the N-terminus, and the LQLPPLERLTLD nuclear export signal at the C-terminus, was reverse translated using the most-preferred *Chlamydomonas* codons. The first *RBCS2* intron was inserted (AG/intron/GG) and the sequence was synthesized with BbsI restriction sites at the 5’- and 3’-termini (producing GACT and AATG overhangs) and cloned into pBS SK+ by GeneCust (Luxembourg), yielding pMBS977. To target APEX2 to the chloroplast, sequences encoding the HSP70B chloroplast transit peptide containing the first *HSP70B* intron, were amplified by PCR on pMBS639 (Niemeyer et al., 2021) (Supplemental Table S1). The resulting 249-bp PCR product, pMBS977, and destination vector pAGM1276 (Weber et al., 2011) were subjected to a restriction-ligation reaction with BbsI, resulting in level 0 vector pMBS454, placing *cp70B-APEX2* into the B2 position. For C-terminal APEX2 fusions, pMBS527 was used as a template for PCR to amplify the APEX2 gene with flanking BbsI restriction sites producing TTCG and GCTT overhangs (Supplemental Table 1). The PCR product and pAGM1301 were subjected to a restriction-ligation reaction with BbsI, resulting in level 0 vector pMBS527, placing *APEX2* into the B5 position.

The BioID (BirA*) amino acid sequence (Choi-Rhee et al., 2004; Roux et al., 2012) with a C-terminal GGGGS-linker was reverse translated using the most-preferred *Chlamydomonas* codons and equipped with the transit peptide sequence of HSP70B containing the first *HSP70B* intron. The fifth *HSP70B* intron was inserted into the BioID coding sequence (AG/intron/GG). Synthesis and cloning into the XbaI-XhoI site of pBS SK+ was done by GeneCust (Luxembourg), yielding pMBS976. Since the original gene design was not compatible with the MoClo syntax, we amplified three fragments by PCR to remove two BsaI sites from the first *HSP70B* intron and the BirA* coding sequence and to introduce flanking BbsI restriction sites giving rise to CCAT and AATG overhangs (primers used and product sizes are listed in Supplemental Table S1). PCR products and destination vector pAGM1276 were subjected to a restriction-ligation reaction with BbsI, resulting in level 0 vector pMBS197.

For C-terminal TurboID fusions, the TurboID protein (Branon et al., 2018), equipped with an N-terminal SGGGG-linker, was reverse translated using the most-preferred *Chlamydomonas* codons and the fifth *HSP70B* intron (AG/intron/GC) was inserted. The sequence was synthesized with flanking BsaI restriction sites (producing TTCG and GCTT overhangs) and cloned into the pUC57 vector by BioCat (Heidelberg, Germany), yielding level 0 construct pMBS512 with *TurboID* in the B5 position. To target TurboID to the chloroplast, the TurboID protein, equipped with a C-terminal GGGGS-linker, was reverse translated using the most-preferred *Chlamydomonas* codons and the fifth *HSP70B* intron (AG/intron/GC) was inserted. The sequence was synthesized with flanking BbsI restriction sites (producing TCAG and AATG overhangs) and cloned into the pUC57 vector by BioCat (Heidelberg, Germany), yielding pMBS513. Sequences encoding the HSP70B chloroplast transit peptide and the first *HSP70B* intron were amplified by PCR on pMBS639 (Supplemental Table S1). The resulting 245-bp PCR product, pMBS513, and destination vector pAGM1276 were subjected to a restriction-ligation reaction with BbsI, resulting in level 0 vector pMBS515, placing *cp70B-TurboID* into the B2 position. All PCRs were done with KAPA HiFi PCR Kit (KapaBiosystems) following the manufacturer’s instructions. Correct cloning of all level 0 constructs made was verified by Sanger sequencing. Level 0 constructs for all baits and biotin activases are shown in Supplemental Figure 1.

*Level 1 and 2 constructs –* The newly constructed level 0 parts were complemented with level 0 parts (pCM) from the *Chlamydomonas* MoClo toolkit (Crozet et al., 2018) to fill the respective positions in level 1 modules as follows: A1-B1 – pCM0-015 (*HSP70A-RBCS2* promoter + 5’ UTR); A1-B2 – pCM0-020 (*HSP70A-RBCS2* promoter + 5’ UTR); B2 – pMBS454 (*chloroplast targeted APEX2*), pMBS197 (*chloroplast targeted BioID),* pMBS515 (*chloroplast targeted TurboID*) and pMBS640 (*CDJ1 chloroplast transit peptide*, Niemeyer et al. (2021)); B3/4 – pMBS375 (*CGE1*), pMBS478 (*VIPP1,* Gupta et al. (2021)), pMBS277 (*VIPP2*, Theis et al. (2020)) or pCM0-067 (*mCherry*); B5 – pCM0-100 (*3xHA*), pCM0-101 (*MultiStop*) or pMBS512 (*TurboID-C*); B6 – pCM0-119 (*RPL23* 3’ UTR). The respective level 0 parts and destination vector pICH47742 (Weber et al., 2011) were combined with BsaI and T4 DNA ligase and directionally assembled into the seven level 1 modules shown in Supplemental Table 2. The level 1 modules were then combined with pCM1-01 (level 1 module with the *aadA* gene conferring resistance to spectinomycin flanked by the *PSAD* promoter and terminator) from the *Chlamydomonas* MoClo kit, with plasmid pICH41744 containing the proper end-linker, and with destination vector pAGM4673 (Weber et al., 2011), digested with BbsI, and ligated to yield seven of the eight level 2 devices displayed in Supplemental Table 2. For the VPL2 construct pMBS970, level 0 parts were directly assembled into level 2 destination vector pMBS807 already containing the *aadA* resistance cassette (Niemeyer and Schroda, 2022). All newly generated level 0 and level 2 plasmids can be ordered from the *Chlamydomonas* Research Center (https://www.chlamycollection.org/).

### Cloning, expression, and purification of recombinant BirA

The BirA coding region was amplified by colony PCR from TOP10F’ cells (Invitrogen) (Supplemental Table 1). The 981-bp PCR product was digested with BamHI and HindIII and cloned into BamHI-HindIII-digested pETDuet-1 vector (Novagen), giving pMS977. BirA was expressed with an N-terminal hexa-histidine (6xHis) tag in *E. coli* Rosetta cells (DE3, Novagen) after inducing expression with 1 mM IPTG for 16 h at 20°C and purified by cobalt-nitrilotriacetic acid affinity chromatography according to the manufacturer’s instructions (G-Biosciences), including a washing step with 5 mM Mg-ATP. Eluted BirA was gel filtrated using an Enrich SEC650 column. The purity of the recombinant protein was analyzed by Coomassie brilliant blue staining (Roth) after separating it on a 12% SDS-polyacrylamide gel. Fractions containing BirA were pooled and concentrated in Amicon® Ultra-4 Centrifugal Filter Units (Ultracel®-3K, Merck Millipore Ltd), with a subsequent buffer exchange to 6 M Urea, 50 mM NaCl, 20 mM Tris-HCl, pH 7.5. The protein concentration was determined by NanoDrop 2000 (ThermoFischer Scientific) based on the molar extinction coefficient and molecular weight of 6xHis-tagged BirA. 1 mg of purified BirA protein was used for the raising of an antiserum in rabbits according to a 3-month standard immunization protocol (Bioscience, bj-diagnostik, Göttingen).

### Protein analyses

Protein extractions, SDS-PAGE, semi-dry blotting and immunodetections were carried out as described previously (Liu et al., 2005; Schulz-Raffelt et al., 2007). Sample amounts loaded were based on protein determination as described by (Bradford, 1976) or based on chlorophyll concentrations (Porra et al., 1989). Antisera used were against BirA (this work), CDJ1 (Willmund et al., 2008), CGE1 (Schroda et al., 2001), CLPB3 (Kreis et al., 2022), DEG1C (Theis et al., 2019b), the HA epitope (Sigma-Aldrich H3663), HSP22E/F (Rütgers et al., 2017), HSP70B (Schroda et al., 1999), HSP90C (Willmund and Schroda, 2005), mCherry (Crozet et al., 2018), RPL1 (Ries *et al.,* 2017), and VIPP1 (Liu et al., 2005). Anti-rabbit-HRP (Sigma-Aldrich) and anti-mouse-HRP (Santa Cruz Biotechnology sc-2031) were used as secondary antibodies. For the detection of biotinylated proteins, proteins were separated on a 12% SDS-PAGE gel and transferred to a nitrocellulose membrane, stained with Ponceau S (1 minute in 0.1% w/v Ponceau S in 5% acetic acid/water), and blocked with “biotin blocking buffer” (3% w/v BSA and 0.1% Tween-20 in PBS) at 22°C for 30 min. The blots were immersed in streptavidin-HRP (1:20,000 dilution, Abcam ab7403) or anti-Biotin-HRP (1:40,000, Sigma-Aldrich A0185) in biotin blocking buffer at 22°C for 60 minutes, then rinsed 3 times with PBS-T for 5 minutes. Detections were performed via enhanced chemiluminescence (ECL) and the FUSION-FX7 Advance™imaging system (PEQLAB) or ECL ChemoStar V90D (INTAS Science Imaging). Densitometric band quantifications after detections were done by the FUSIONCapt Advance program (PEQLAB).

### *In vivo* labeling with biotin-phenol

Cells were preincubated with biotin-phenol for time periods of 10 min up to 24 h at 22°C. From a 100 mM biotin-phenol stock in dimethyl sulfoxide (DMSO), biotin-phenol was diluted directly into cell cultures to a final concentration of 500 µM. Labeling was started by the addition of H_2_O_2_ to a final concentration of 1 mM and allowed to proceed for time periods of 1 min up to 1 h. Labeling was stopped by transferring cells on ice and washing with fresh, ice-cold TAP medium.

### *In vitro* labeling with biotin-phenol

10 mL of exponentially growing cells were harvested by centrifugation for 3 min at 4,000 g and 22°C, resuspended in 1 mL KMH buffer (20 mM HEPES-KOH pH 7.2, 10 mM KCl, 1 mM MgCl_2_, 154 mM NaCl, 1x cOmplete^TM^, EDTA-free protease inhibitor cocktail (Roche)). Cells were lysed by 3 freezing and thawing cycles. Soluble proteins were prepared by centrifugation at 20,000 g for 30 min at 4°C. Biotin-phenol was then added from a 500 mM stock in DMSO to a final concentration of 500 µM. After a preincubation period of 30 min at 22°C, 1 mM H_2_O_2_ was added for 1 min. Labeling was stopped by transferring the proteins on ice.

### *In vivo* biotin-labeling without biotin addition and streptavidin affinity purification

500 mL of exponentially growing cells were harvested by centrifugation for 2 min at 4000 g and 22°C and resuspended in 15 mL RIPA buffer (50 mM Tris-HCl pH 7.5, 150 mM NaCl, 0.1% SDS, 0.5% sodium deoxycholate, 1% Triton X-100, 2.5 mM EDTA, 1 mM DTT, 1× protease inhibitor cocktail (Roche), 1 mM phenylmethylsulfonyl fluoride (PMSF)). Cells were snap-frozen in liquid nitrogen and stored at -80°C prior to cell lysis. Cell samples were vortexed, sonicated (4 x 10 s, minimal output, 7 cycles with 1-min breaks on ice) and centrifuged at 14,000 x g and 4°C for 30 min. Precleared whole-cell lysates were next applied to streptavidin agarose resin (200 µL, Thermo-Fischer), and incubated overnight at 4°C on a SB2 rotator (Stuart). The resin was then washed at least twice in 10 mL RIPA buffer, once in 1 mL 1 M KCl, once in 1 mL 0.1 M Na_2_CO_3_ at 4°C. After transfer to a fresh tube at 22°C, the resin was washed once in urea buffer (2 M urea in 10 mM Tris-HCl, pH 8.0), twice with RIPA buffer, followed by a further transfer to a fresh tube. Proteins were eluted by boiling the resin for 10 min at 98°C in elution buffer (125 mM Tris pH 7.5, 4% SDS, 20 mM DTT, 2 mM biotin, 0.05% bromphenol blue) and subjected to SDS-PAGE for immunoblotting as well as to mass spectrometry analysis.

### *In vivo* biotin-labeling with biotin addition and streptavidin affinity purification

Biotin was added from a 100 mM stock in DMSO to a final concentration of 1 mM into a 250-mL culture of exponentially growing cells for 4 h. If indicated, H_2_O_2_ treatment was done in parallel. Labeling was stopped by transferring the cells to ice and washing once with ice-cold fresh TAP medium to remove the biotin. Cells were harvested by centrifugation for 2 min at 4000 g at 4°C, resuspended in 6 mL RIPA buffer, snap-frozen in liquid nitrogen and stored at -80°C prior to cell lysis. The cells were thawed on ice, vortexed, sonicated (4 x 10 s, minimal output, 6x cycles with 30 s breaks on ice), and centrifuged at 20,000 x g for 20 min at 4°C. Precleared cell lysates were applied to streptavidin agarose resin (100 µl, Thermo), and incubated overnight at 4°C on a rotator. Washing and elution were performed as described above.

### *In vitro* biotin-labeling on isolated membrane proteins and streptavidin affinity purification

500 mL of exponentially growing cells were harvested by centrifugation for 2 min at 4000 g at 22°C, resuspended in 5 mL KMH buffer, snap-frozen in liquid nitrogen, and stored at -80°C prior to cell lysis. The cells were broken by three freeze/thaw cycles, distributed to 2 ml tubes to separate soluble and membrane fractions by centrifugation at 10,000 g for 30 min and 4°C. The soluble fraction was discarded. Membranes were homogenized using a potter in 1.5 ml KMH buffer supplemented with 1 mM PMSF. Biotin was added to the lysate from a 500 mM biotin stock dissolved in H_2_O to a final concentration of 500 µM and an ATP regeneration system was added (2.5 mM ATP, 80 mM phosphocreatine disodium salt hydrate, 0.125 µg/µl creatine phosphokinase from bovine heart). Labeling was stopped after 30 min by transferring the cells to ice and the addition of extraction reagents (0.1% SDS, 0.5% sodium deoxycholate, 1% Triton X-100, 2.5 mM EDTA, 1 mM DTT and 1 × protease inhibitor cocktail (Roche)). The samples were sonicated (4 x 30s, minimal output, 6 cycles with 1.5-min breaks on ice), centrifuged at 20,000 g for 20 min, followed by the removal of biotin by desalting on PD10 columns (GE Healthcare). The desalted lysates were then applied to streptavidin agarose resin (100 µl, Thermo), and incubated overnight at 4°C on a rotator. Further processing was done as described for *in vivo*-labelling above.

### MS sample preparation and mass spectrometry

For in-gel digestion, biotinylated proteins were precipitated overnight at -20°C after adding ice-cold acetone to a final volume of 80%. Precipitated proteins were pelleted by centrifugation for 20 min at 25,000 g and 4°C. After washing with 80% acetone, the pelleted proteins were air-dried, resuspended in Laemmli buffer (Laemmli, 1970) and allowed to just migrate into the separating gel of a 10% SDS-polyacrylamide gel and stained with colloidal Coomassie G (Candiano et al., 2004). Entire protein lanes were excised in 2-3 bands to separate streptavidin, cut in cubes and destained (40% methanol, 7% acetic acid). Alkylation of cysteines and tryptic digest over night at 37°C was performed as described earlier (Veyel et al., 2014). Hydrophilic peptides were extracted from the gel pieces with 10% acetonitrile and 2% formic acid for 20 min and afterwards all other tryptic peptides were extracted with 60% acetonitrile, 1% formic acid. Samples were then desalted according to Rappsilber et al. (2007). Mass spectrometry on a LC-MS/MS system (Eksigent nanoLC 425 coupled to a TripleTOF 6600, ABSciex) was performed basically as described previously (Hammel et al., 2018). For peptide separation, a HPLC flow rate of 4 μl/min was used and gradients (buffer A 2% acetonitrile, 0.1% formic acid; buffer B 90% acetonitrile, 0.1% formic acid) ramped within 48 min from 2% to 35% buffer B, then to 50% buffer B within 5 min and finishing with wash and equilibration steps. MS^1^ spectra (350 m/z to 1250 m/z) were recorded for 250 ms and 35 MS/MS scans (110 m/z to 1600 m/z) were triggered in high sensitivity mode with a dwell time of 60 ms resulting in a total cycle time of 2400 ms. Analysed precursors were excluded for 10 s, singly charged precursors or precursors with a response below 150 cps were excluded completely from MS/MS analysis.

### Evaluation of MS data

The analysis of MS runs was performed using MaxQuant version 1.6.0.1 (Cox and Mann, 2008). Library generation for peptide spectrum matching was based on *Chlamydomonas reinhardtii* genome release 5.5 (Merchant et al., 2007) including chloroplast and mitochondrial proteins as well as TurboID and mCherry. Oxidation of methionine and acetylation of the N-terminus were considered as peptide modifications. Maximal missed cleavages were set to 3 and peptide length to 6 amino acids, the maximal mass to 6000 Da. Thresholds for peptide spectrum matching and protein identification were set by a false discovery rate (FDR) of 0.01. The mass spectrometry proteomics data have been deposited to the ProteomeXchange Consortium via the PRIDE partner repository with the dataset identifier PXDxxxxxx (Perez-Riverol et al., 2019).

### MS data analysis – identification of enriched proteins

Filtering and statistical analysis were done with Perseus (version 2.0.3.0.) (Tyanova et al., 2016). The ‘proteinGroups.txt’ output file from MaxQuant was imported into Perseus using the intensity values as main category. The data matrix was filtered to remove proteins marked as ‘only identified by site’, ‘reverse’ and ‘potential contaminant’. Intensity values were log2 transformed and proteins were removed that were not identified/quantified in all three replicates of at least one strain under one condition. Normalization was achieved by subtracting the median using columns as matrix access. Next, missing values were imputed for statistical analysis using the ‘Replace missing values from normal distribution’ function with the following settings: width = 0.3, down shift = 1.8, and mode = total matrix. To identify proteins enriched in T_ID_-expressing samples versus the WT control, unpaired two-tailed Student’s t-tests were performed comparing the T_ID_CGE1, VIPP1T_ID_, VIPP2T_ID_, VPL2T_ID_ or mCherryT_ID_ with the corresponding WT samples at each time point. The integrated modified permutation-based FDR was used for multiple sample correction with an FDR of 0.05, an S0 of 1, and 250 randomizations to determine the cutoff. Significantly enriched proteins were kept. To further identify proteins that are significantly enriched in T_ID_CGE1, VIPP1T_ID_, VIPP2T_ID_, and VPL2T_ID_ versus mCherryT_ID_, corresponding t-tests were performed on the reduced datasets using the same parameters as for the first t-tests. Finally, to determine proteins that were enriched in at least one of the two treatments (CL or HS, CL or H_2_O_2_) compared to mCherryT_ID_, the cutoff parameters were set to a q-value <0.05 and minimal fold change log2 ≥ 1. Scatterplots were made in Excel using t-test results exported from Perseus.

## Supplemental Files

**Supplemental Figure S1.** Level 0 constructs for baits and biotin activators used in this study.

**Supplemental Figure S2.** Screening for transformants accumulating CGE1 and mCherry with N-terminal fusions to APEX2.

**Supplemental Figure S3.** Dependency of APEX2 activity on biotin-phenol preincubation time and labeling reaction time.

**Supplemental Figure S4.** Screening for transformants accumulating CGE1 N-terminally fused to BioID.

**Supplemental Figure S5.** Screening for transformants accumulating TurboID fusions to VIPP1, VIPP2, CGE1, and mCherry.

**Supplemental Figure S6.** Production of recombinant BirA and characterization of the antiserum raised against it.

**Supplemental Figure S7.** Impact of TurboID mediated biotinylation on cell fitness.

**Supplemental Figure S8.** Comparison of structures and sequences of CGE2 and CGE1.

**Supplemental Figure S9.** Structures and alignments of VPL2 orthologs.

**Supplemental Figure S10.** Screening of transformants accumulating VPL2 with a C-terminal TurboID fusion.

**Supplemental Table S1.** Primers used for cloning.

**Supplemental Table S2.** MoClo constructs employed and generated.

**Supplemental Table S3.** Predicted molecular masses of fusion proteins.

**Supplemental Data Set S1.** Proteins significantly enriched after *in vivo* biotinylation in TurboID-CGE1 vs. WT and in TurboID-CGE1 vs. mCherry-TurboID.

**Supplemental Data Set S2.** Proteins significantly enriched after *in vivo* biotinylation in VIPPs-TurboID vs. WT and in VIPPs-TurboID vs. mCherry-TurboID.

**Supplemental Data Set S3.** Proteins significantly enriched after boosted *in vitro* biotinylation on isolated membranes in VIPPs-TurboID vs. WT and in VIPPs-TurboID vs. mCherry-TurboID.

**Supplemental Data Set S4.** Proteins significantly enriched after boosted *in vivo* biotinylation in VIPPs-TurboID vs. WT and in VIPPs-TurboID vs. mCherry-TurboID.

**Supplemental Data Set S5.** Proteins significantly enriched after boosted *in vivo* biotinylation in VPL2-TurboID vs. WT and in VPL2-TurboID vs. mCherry-TurboID.

**Supplemental Data Set S6.** Cellular localization of proteins identified in the TurboID-mediated VIPP1/2, VPL2, and CGE1 proxiomes.

## Acknowledgements

This work was supported by the Deutsche Forschungsgemeinschaft [SFB/TRR175, project C02; SPP1927, project Schr 617/11-1].

## Author Contributions

E.K. and K.K. generated all constructs and performed all experiments. F.S performed the mass spectrometry analyses. M.S. conceived and supervised the project and wrote the article with contributions from all authors.

